# Direct effects on gene transcription revealed by global and CRISPR-targeted oxidative DNA damage

**DOI:** 10.64898/2026.10.26.761859

**Authors:** Luana Cintori, Catherine Chailleux, Pauline Maller, Anne-Marie Di Guilmi, Capucine Siberchicot, Tomas Caetano, Emma Lehir, Virginie Jouffret, Celine Reyes, Marion Aguirrebengoa, Anna Campalans, Valérie Bergoglio, Didier Trouche, Yvan Canitrot

**Affiliations:** Chromatin and proliferation team, University of Toulouse, CNRS, Center for Integrative Biology (CBI),169 avenue Marianne Grunberg-Manago, 31400, Toulouse, France; Université Paris-Saclay, Université Paris Cité, Inserm, CEA, Institut de Biologie François Jacob, UMR Genetic Stability, Stem Cells and Radiation, F-92260 Fontenay-aux-Roses, France; BigA facility, University of Toulouse, CNRS, Center for Integrative Biology (CBI), Toulouse, France

**Author notes:** These authors contributed equally to this work (Valérie Bergoglio, Didier Trouche, Yvan Canitrot).

## Abstract

Oxidative stress induces rapid expression changes of specific genes. However, whether these changes are due to direct and local DNA oxidation and whether they depend on the repair machinery is still debated. Here, we show by total RNA-Seq that upon oxidative stress, many genes are either rapidly repressed or activated. The early transcriptional response also includes the activation of many read-through RNAs. Importantly, degradation of the endogenous OGG1 DNA glycosylase showed that the early repression of these genes partially depends on OGG1, therefore demonstrating that these genome expression changes are due to the processing of oxidized DNA. We further demonstrate by ChIPseq against total or S5-phosphorylated RNA polymerase II that transcriptional regulation by oxidative stress involves changes in the recruitment of RNA polymerase II to transcription start sites. Finally, through the use of a chemoptogenetic system allowing the targeting of ROS to chosen genomic loci, we were able to reproduce repression of some genes through the local induction of oxidative DNA damage at their promoters, demonstrating that local DNA oxidation directly affects transcription of genes. Altogether, our results show that local oxidative DNA damage leads to alterations in the activity of specific promoters by a mechanism dependent on repair processes.

## Introduction

The genomes of eukaryotic cells are under constant threat from a variety of damaging agents. To counteract these threats, cells have evolved multiple DNA repair pathways that mitigate the deleterious consequences of unrepaired DNA lesions. Among the most significant sources of such damage is oxidative stress, a condition arising from an imbalance between the production and detoxification of reactive oxygen species (ROS). ROS, which originate from both endogenous and exogenous sources, can oxidize a wide range of cellular components, including DNA ^1^. DNA oxidation represents the most prevalent form of DNA damage and is a major driver of mutagenesis, as well as a contributing factor to cancer, aging, and neurodegenerative diseases ^2^. ROS induce a diverse spectrum of DNA lesions, with oxidized bases primarily repaired through the Base Excision Repair (BER) pathway. This pathway is initiated by DNA glycosylases, which recognize and excise damaged bases, generating abasic sites. These abasic sites are subsequently converted into single-strand breaks (SSBs) by the apurinic/apyrimidinic endonuclease (APE1), and the resulting gaps are filled by DNA polymerase beta and sealed by DNA ligase III ^3^. Among these glycosylases, OGG1 specifically targets the most abundant oxidized base, 8-oxo-guanine (8-oxo-G), as guanine is the most oxidation-prone nucleotide in DNA ^4^.

The impact of oxidative stress on DNA function, particularly on transcription has been the subject of extensive investigation but is still a matter of debate. Some data show that oxidative stress triggers a widespread repression of transcriptional activity ^5–7^. However, unlike bulky UV-induced lesions, which block transcription elongation and require repair via transcription-coupled nucleotide excision repair (TC-NER), oxidative DNA lesions are typically non-bulky and can be bypassed by RNA polymerase II, as demonstrated *in vitro* ^8^. Consistent with this, some studies based on using genome-wide techniques observed both repression and activation of gene transcription ^9,10^ upon global oxidative stress induction. Furthermore, these conditions promote the transcription of read-through RNAs ^11^, ie RNAs produced downstream of genes and which could play a role in euchromatin protection against stress ^11^.

Investigations of specific promoters have further elucidated how oxidative stress modulates gene expression. In particular, 8-oxo-G can occasionally function as an epigenetic-like mark (for review ^12^). For example, the oxidative stress-induced activation of the *VEGF* gene has been linked to the accumulation of 8-oxo-G at its promoter, where these lesions may facilitate the formation of G-quadruplex structures ^13^. However, many of these findings rely on global oxidative stress induction, or were performed only *in vitro* ^8,14^ , which complicates the interpretation of results due to potential indirect effects. For instance, changes in genome-wide gene expression could stem from the activation of cellular checkpoints or signaling pathways rather than from localized DNA oxidation. Additionally, the role of OGG1 has largely been inferred from comparisons between wild-type (WT) and OGG1-knockout (KO) cells. This approach may introduce confounding variables, such as compensatory mechanisms arising from redundancy among DNA glycosylases in KO cells.

To overcome these limitations, we developed two key innovations. First, we engineered a cell line enabling the rapid and efficient degradation of OGG1. Second, we adapted an optogenetic system to generate ROS production around chosen DNA regions, allowing precise spatiotemporal control of oxidative damage. Using genome-wide analyses and this targeted approach, we demonstrate that local oxidative DNA damage and its processing by OGG1 modulate the transcription of specific genes through alterations in RNA polymerase II (Pol II) recruitment at transcription start sites (TSSs).

## Results

### Generation of a cell line allowing the rapid degradation of OGG1 DNA glycosylase

To investigate the role of OGG1 DNA glycosylase in the transcriptional response to oxidative DNA damage, we generated a U2OS osteosarcoma cell line enabling the rapid and inducible degradation of OGG1. Previous studies on transcriptional function of OGG1 have primarily relied on OGG1 KO cells ^9,15^ , which may allow cellular adaptation to the absence of OGG1, or on OGG1 inhibitors, which can have off-target effects ^16^.

To overcome these limitations, we used genome editing to tag endogenous OGG1 with an FKBP12^F36V^ inducible degron sequence and an HA tag at both alleles in U2OS cells (Figure 1A). The FKBP12 sequence was added at the C-terminus of OGG1, as a GFP tag at this position was previously shown not to affect OGG1 activity and chromatin localization following oxidative DNA damage induction ^17,18^. Treatment with the dTAGV1 molecule led to rapid and efficient degradation of the OGG1-FKBP-HA protein, as observed by Western blot (Figure 1B and Figure Supp 1A). Importantly, we confirmed that the OGG1-FKBP12-HA fusion protein retains glycosylase activity that is completely lost upon degradation of OGG1 indicating that no backup pathway allowing the processing of 8oxoG was present in this condition (Figure 1C and Figure Supp 1B). This cell line thus provides a powerful model to study OGG1 function without the confounding effects of cellular adaptation.

**Figure 1.**
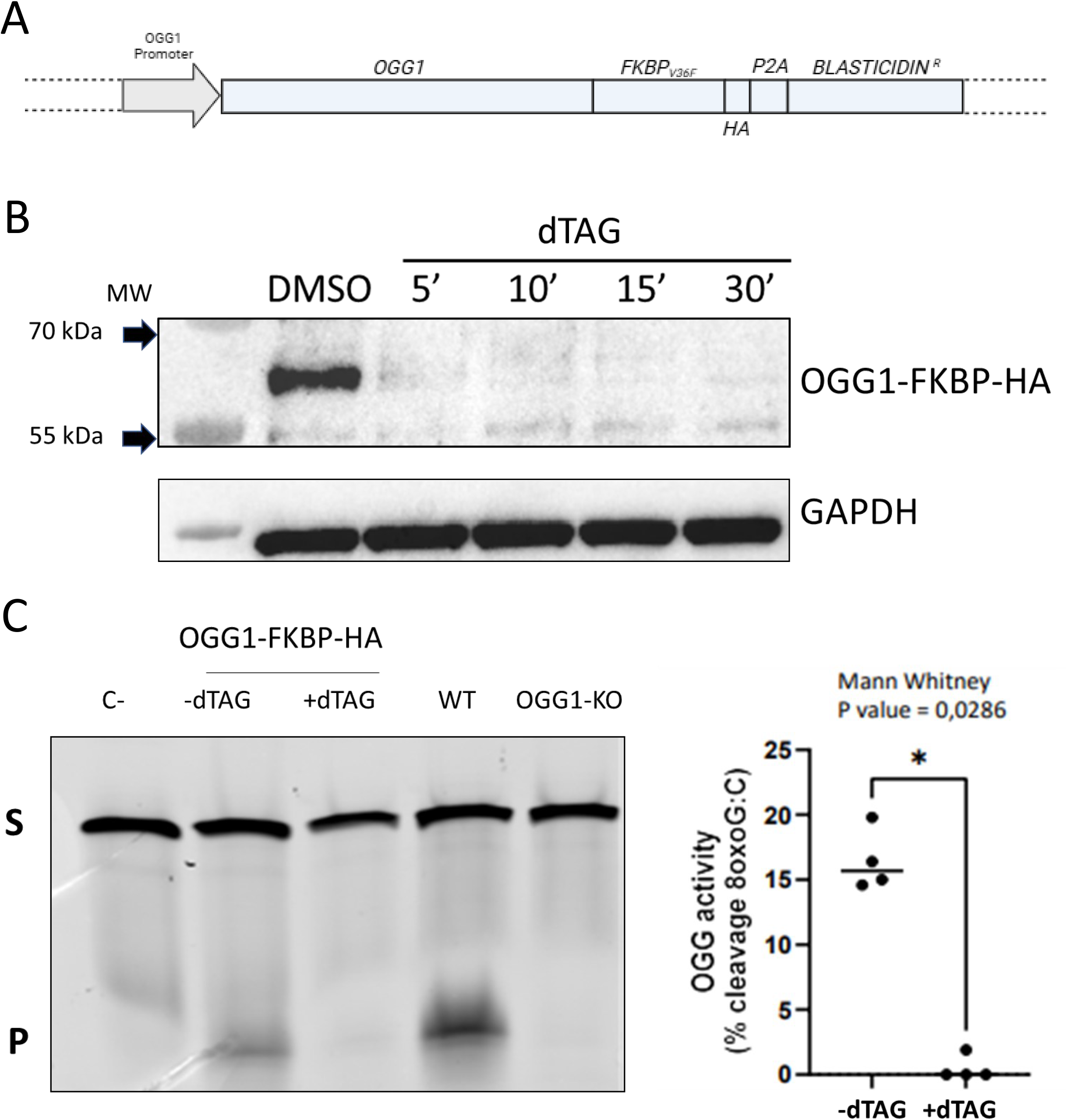
Generation of the OGG1-FKBP-HA cell line. **A**. Scheme of genetic modification created in the U2OS cell line. **B**. Effect of cell treatment with dTAGV1 on OGG1 expression examined by Western blot. Cells were treated with dTAGV1 for the indicated time and total cell extracts prepared and tested by Western blot using anti-HA and anti-GAPDH as a loading control. A representative experiment out of 2 is shown. **C.** Checking of the enzymatic activity of OGG1-FKBP-HA construct. Activity gel measuring 8-oxoG incision was performed with cell extracts from WT, KO or OGG1-FKBP-HA U2OS cell extracts treated or not with dTAGV1 for 1h. A typical experiment is shown, and a quantification of 4 independent experiments is shown on the right. The star (*) indicate a p-value < 0.05 (Mann-Whitney).

### Oxidative stress treatment results in rapid genome expression changes

To identify genome-wide expression changes induced by acute oxidative stress, we treated U2OS-OGG1-FKBP-HA cells with or without dTAG to deplete OGG1, and with or without the ROS-inducing agent KBrO₃ for 30 minutes. In order to investigate the role of OGG1 we used KBrO3 as inducer of oxidative stress for its ability to generate a high proportion of 8-oxo-G ^19,20^. Importantly, such KBrO₃ treatment does not induce a major general decrease of transcription as measured by metabolic labelling (Figure S2). Indeed, after subtracting the background, the median signal decreased by 20% whereas it decreased more than three-fold following H_2_O_2_ treatment as described in ^5–7^. After washing out KBrO₃, cells were harvested after 0, 15 and 60 minutes of recovery (see Figure 2A for the experimental scheme). Total RNA was then prepared and analyzed by RNA-Seq.

**Figure 2.**
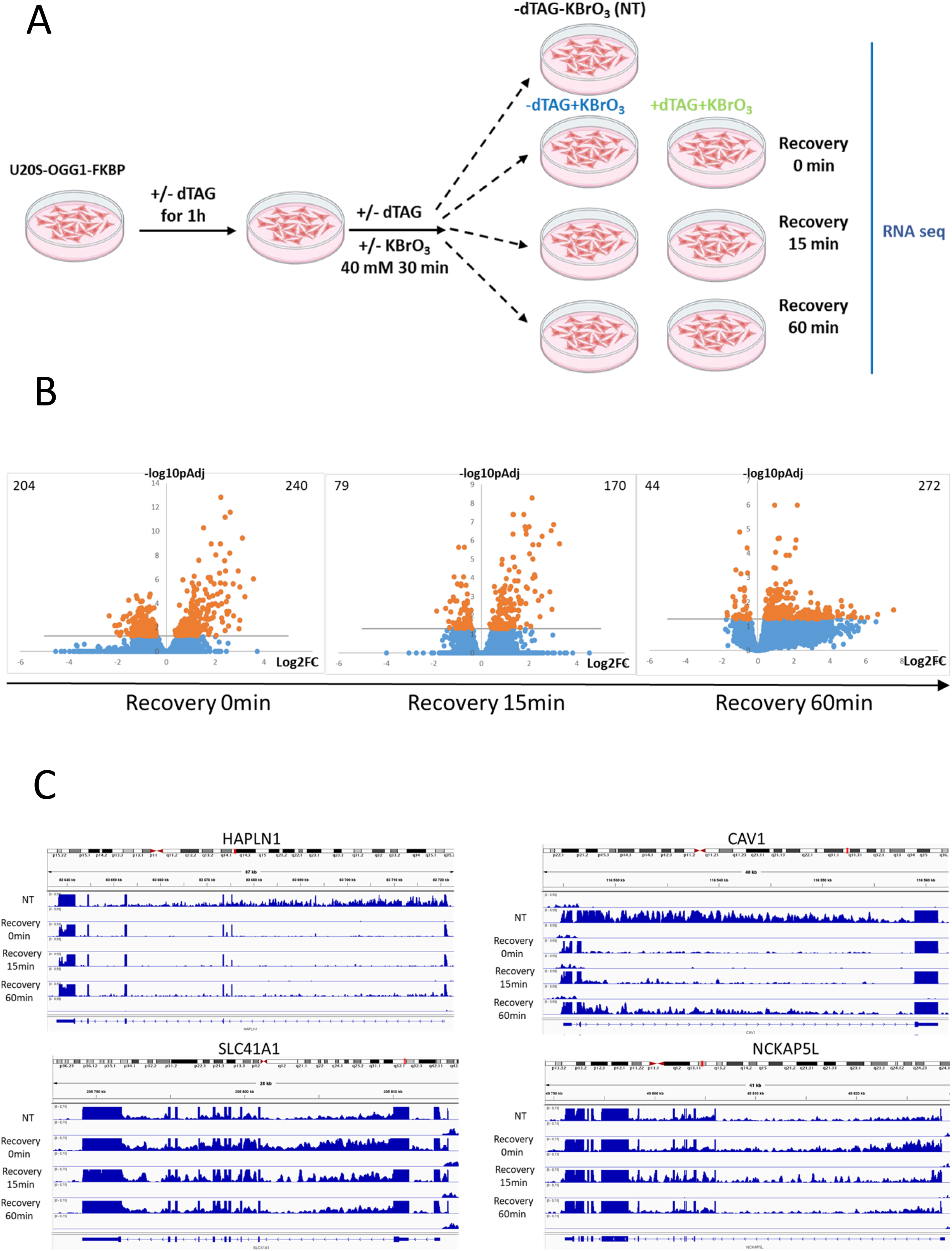
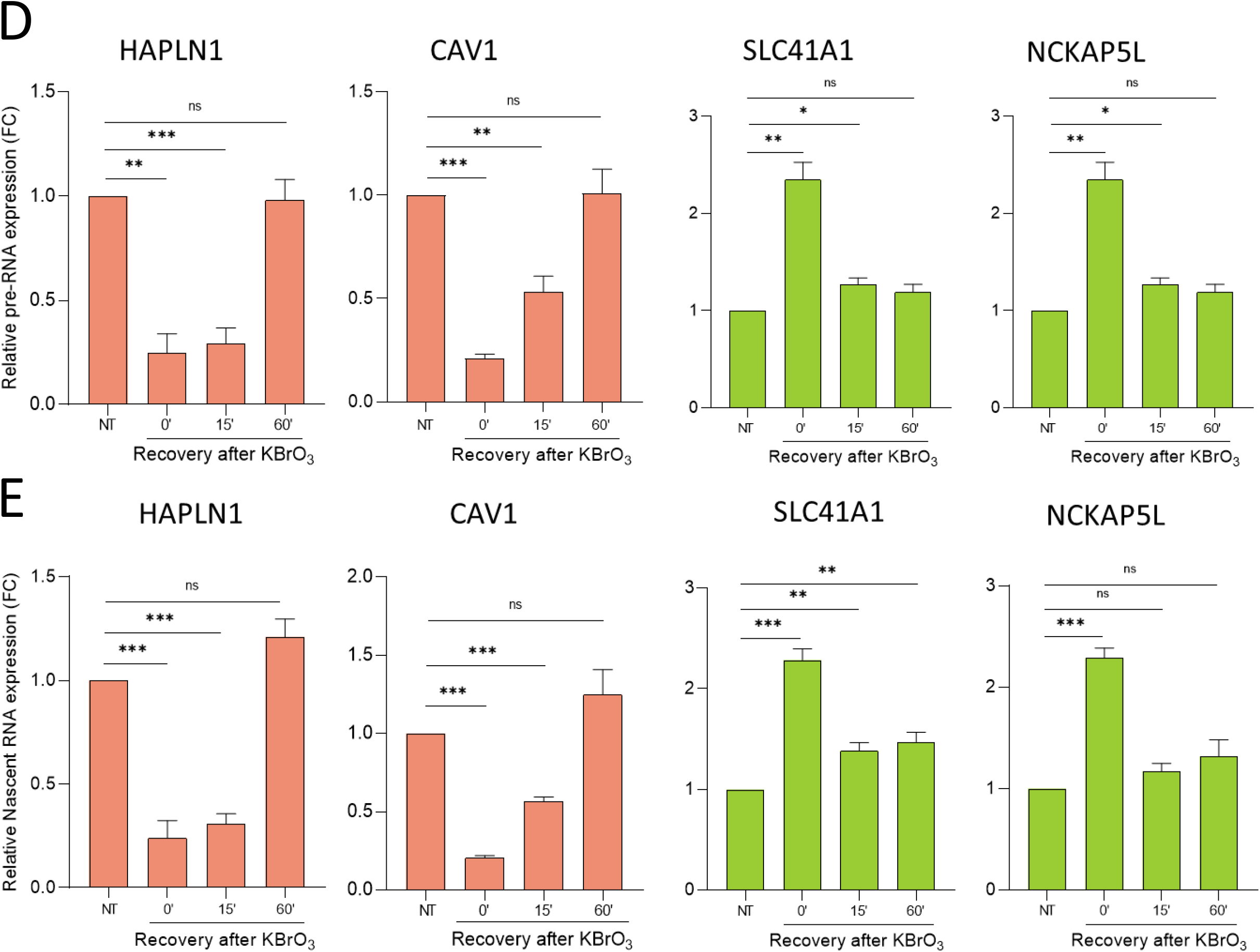
Global oxidative DNA damage leads to rapid transcriptional changes. **A**. Schematic representation of the experimental setting: OGG1-FKBP-HA cells were treated or not with dTAGV1 for 1h then treated or not with KBrO3 for 30 minutes. KBrO3 was then washed, and cells were harvested after the indicated time. **B**. OGG1-FKBP-HA cells were treated or not with KBrO3 for 30 minutes and released for the indicated time. Total RNA was extracted from three independent experiments and subjected to RNA-Seq. The number of intronic reads was then calculated for each gene, and a DGE analysis was performed. Volcano plot showing differential gene expression in the indicated samples compared to the untreated control (-dTAG and no KBrO3) are shown. On the upper corner left and right are the number of genes with -log10pAdj>1,3 and Log2FC<-0.7 or >+0.7. **C**. IGV profiles of a typical experiment representing two repressed genes (HAPLN1 and CAV1) and two activated (SLC41A1 and NCKAP5L) in the minus dTAG conditions. **D**. OGG1-FKBP-HA cells were treated or not with KBrO3 for 30 minutes and released for the indicated time. Total RNA was extracted and analysed by RT-qPCR for CAV1, HAPLN1, NCKAP5L, SLC41A1 as well as the house-keeping RPLP0 premRNAs expression. Results were normalized to RPLP0 expression and standardized relative to 1 in untreated cells. The mean and SDOM from three independent experiments are shown. The stars indicate a p-value (<0.05 for *, <0.01 for ** and <0.001 for ***) (Mann-Whitney test) **E**. OGG1-FKBP-HA cells were treated or not with KBrO3 for 30 minutes and released for the indicated time. Nascent RNA was analysed by RT-qPCR for CAV1, HAPLN1, NCKAP5L, SLC41A1 as well as house-keeping RPLP0 RNAs expression. Results were normalized to RPLP0 expression and standardized relative to 1 in untreated cells. The mean and SDOM from three independent experiments are shown.

We first focused on the response to oxidative stress induction in the absence of dTAG treatment. To identify transcriptional changes, we monitored pre-mRNA expression by analyzing gene intron levels. Due to the very short half-life of introns, their levels largely reflect active gene transcription. Differential expression analysis of pre-mRNA revealed hundreds of genes significantly affected by KBrO₃ treatment immediately after oxidative stress (0 min) or after 15 or 60 minutes of recovery (see the volcano plots in Figure 2B and examples of regulated genes in Figure 2C). Changes observed immediately after oxidative stress seems to be only transient, coming back close to pretreatment levels after 60 minutes of recovery (Figure 2C). Using RT-qPCR on introns, we confirmed the rapid and transient activation of *SLC41A1* and *NCKAP5L* and repression of *HAPLN1* and *CAV1* pre-mRNA expression (Figure 2D). Importantly, analysis of nascent RNAs confirmed that these changes are transcriptional, and not a consequence of changes in pre-mRNA half-life (Figure 2E).

To obtain an integrated view of the dynamic transcriptional response to oxidative stress, we performed unsupervised clustering of differentially expressed genes based on their log₂ fold change (considering genes with an adjusted p value <0.05 and a |log₂FC| > 0.7 in at least one condition). Excluding outliers, this clustering identified four groups of genes in OGG1-expressing cells (Figure 3A). Two groups were characterized by their rapid activation upon oxidative DNA damage (clusters 1 and 2), one group was rapidly repressed (cluster 3, hereafter referred to as "rapidly repressed genes"), and one group was activated only after 1 hour of recovery (cluster 4, hereafter referred to as "late-activated genes") (Figure 3A). Box plot analyses of these populations confirmed these features (Figure 3B). Furthermore, the rapidly-responding genes (clusters 1, 2, and 3) gradually recovered their baseline expression levels during the recovery period following KBrO₃ removal, as already observed in Figure 2 for selected genes. Thus, in response to oxidative stress, we observed three features of the transcriptional response: first, rapid repression or activation of hundreds of genes; second, the gradual recovery of baseline expression levels for these genes; and third, activation of some genes at later time points.

**Figure 3.**
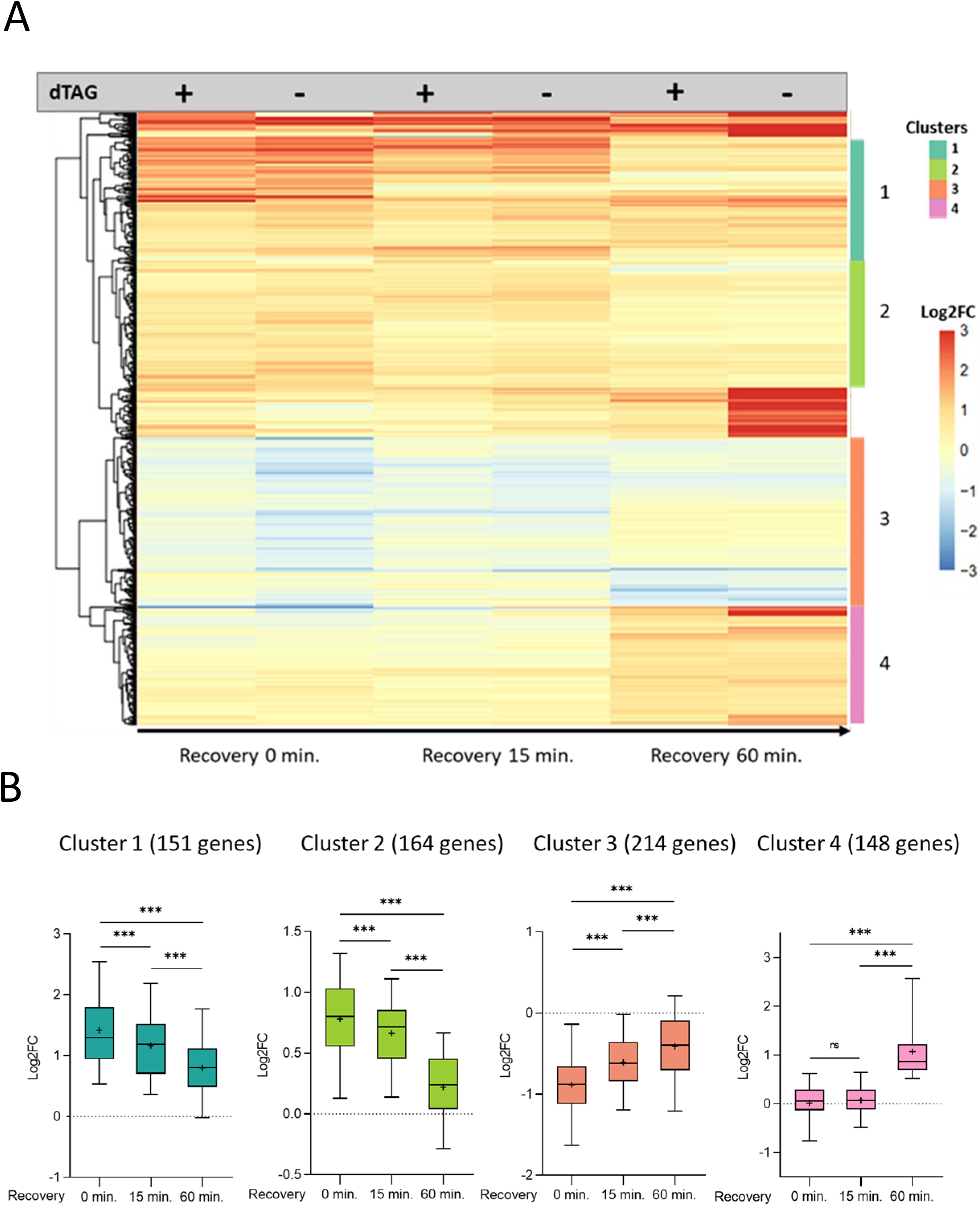
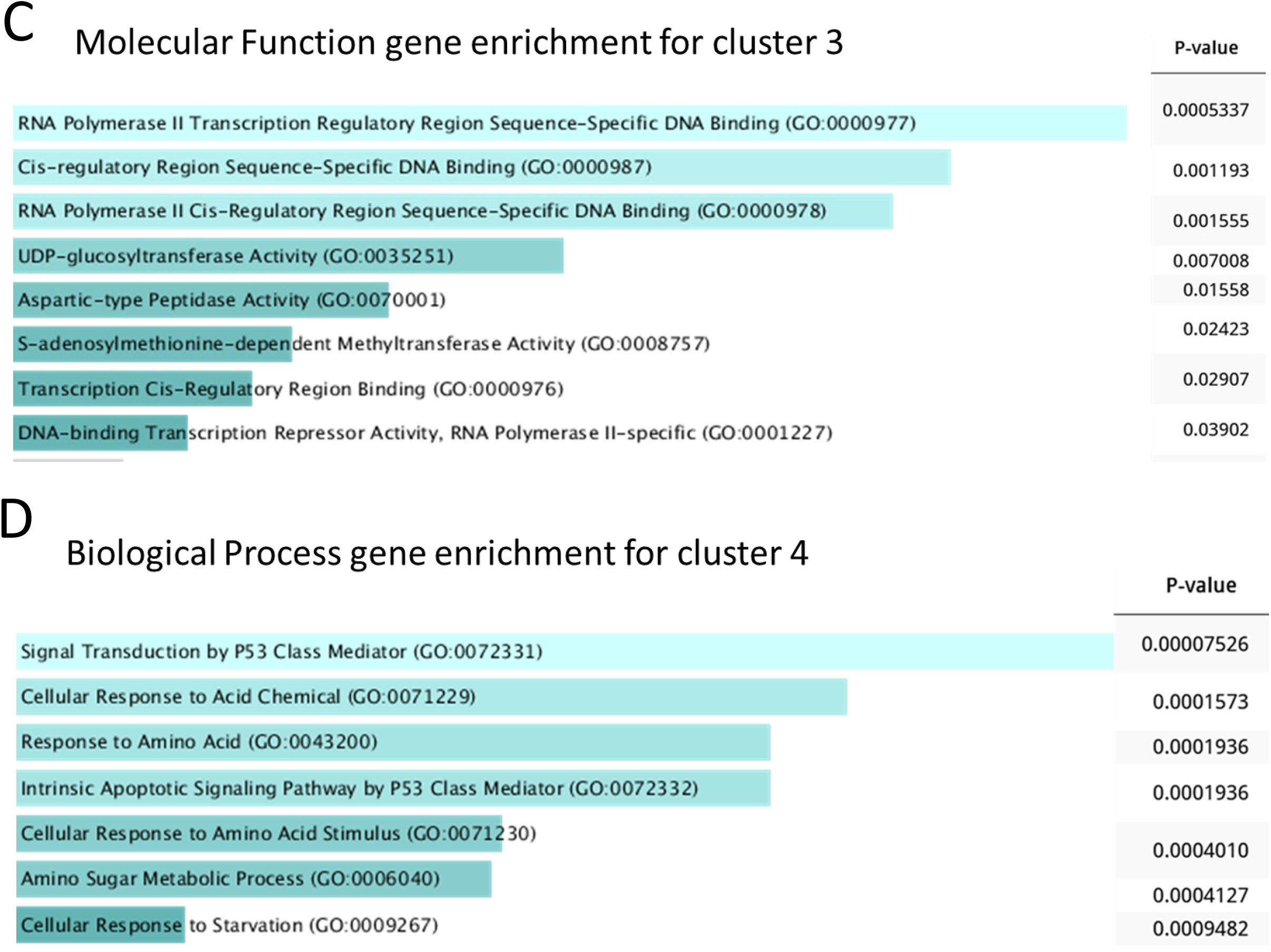
Identification of four clusters of similarly regulated genes. **A**. Genes with a padj<0.05 and a Log2FC<-0.7 or >0.7 in at least one condition from the DGE analysis described in Figure 2B were subjected to unsupervised clustering, leading to the identification of 4 main clusters as indicated by color codes. **B**. Box plots with median, 25 and 75% quantiles showing the evolution of genes from these four clusters during recovery following KBrO3 withdrawal. The stars indicate a p-value (<0.05 for *, <0.01 for ** and <0.001 for ***) (Wilcoxon paired test) **C-D**. GO enrichment analysis for Molecular function performed on cluster 3 (C) and Biological process performed on cluster 4 using Enrichr. The length of the bar represents the significance of that specific gene-set or term. In addition, the brighter the color, the more significant that term is.

Of note, we notice an enrichment of non-coding genes amongst rapidly affected genes, as already observed by others ^10^, which could thus suggest a functional importance of non-coding RNAs in the early response to oxidative DNA damage. Gene ontology (GO) analysis of the Molecular Function category for early-responsive genes, particularly repressed genes, revealed significant enrichment in transcriptional regulation-related genes (Figure 3C and Figure S3). This suggests their role in establishing a global genetic response to oxidative stress. In contrast, late-activated genes did not show enrichment in the same Molecular Function pathways. Instead, they were enriched in Biological Process genes associated with stress-induced signaling pathways (Figure 3D and S3), likely reflecting their activation as a response to oxidative stress.

### Oxidative stress generates read-through RNAs activation

Oxidative stress has been shown to induce transcriptional read-through, i.e. transcription downstream of genes beyond the termination zone, leading to the productions of so-called DoGs RNAs ^11^. We therefore analyzed read-through RNAs expression in our RNA-Seq data. To this end, we first identified read-through RNAs expressed in at least one of our RNA-Seq samples using a pipeline we recently developed (Jouffret et al., unpublished data). This analysis led to the identification of 13316 read-through RNAs. Differential gene expression (DGE) analysis indicated that 1369 read-through RNAs are induced immediately following oxidative stress (see the NF2 and TPT1 genes as examples in Figure 4A and volcano plots in Figure 4B), consistent with published data ^11^. Activation of read-through RNAs originating from the NF2 and TPT1 genes was confirmed on independent experiments by RT-qPCR amplifying a region near TSS (proximal read-through) or farther from TSS (distal read-through) (Figure 4C). Further analysis showed that read-trough RNAs induction decreases during recovery (Figure 4D). Interestingly, a gene ontology performed using the list of genes presenting read-through RNAs shows an enrichment of genes involved in oxidative stress response (Figure 4E), pointing to the functional importance of these read-through RNAs in the response to oxidative stress. Importantly, when examining IGV profiles, we observed that many genes classified as rapidly activated upon oxidative stress were covered by a read-through RNA originating from an upstream gene in the same orientation and activated by oxidative stress (see Figure 4F for two examples). In addition, these genes did not exhibit classical exonic peaks. This result suggests that among rapidly activated genes, some are transcribed from their own promoters, while others are transcribed as part of a read-through RNA from an upstream gene. This is reminiscent to HSV-1 infection, in which observed activation of many genes was found to be due to activation of read-through transcription originating from upstream genes ^21,22^. Upon closer inspection of the two groups of rapidly activated genes, we found that most genes in cluster 1 were covered by a read-through RNA originating from an upstream gene, whereas genes in cluster 2 were primarily activated from their own promoters. To confirm this observation, we performed in untreated cells ChIP-seq for RNA polymerase II phosphorylated on Ser5 (S5-P), which marks transcription initiation events ^23^. Metaprofiles representing S5-P Pol II occupancy around transcription start sites (TSSs) and drawn from two independent replicates showed that cluster 1 genes had very low levels of S5-P at their promoters compared to all genes and to genes from any other cluster (Figure 4G), indicating that the promoters driving cluster 1 genes were inactive. This indicate that the activation of cluster 1 genes is not due to activation of their promoters but rather reflects the induction of a Read-Through RNA activated by oxidative stress. Thus, we excluded cluster 1 genes from further analysis and hereafter refer to cluster 2 genes as "rapidly activated genes."

**Figure 4.**
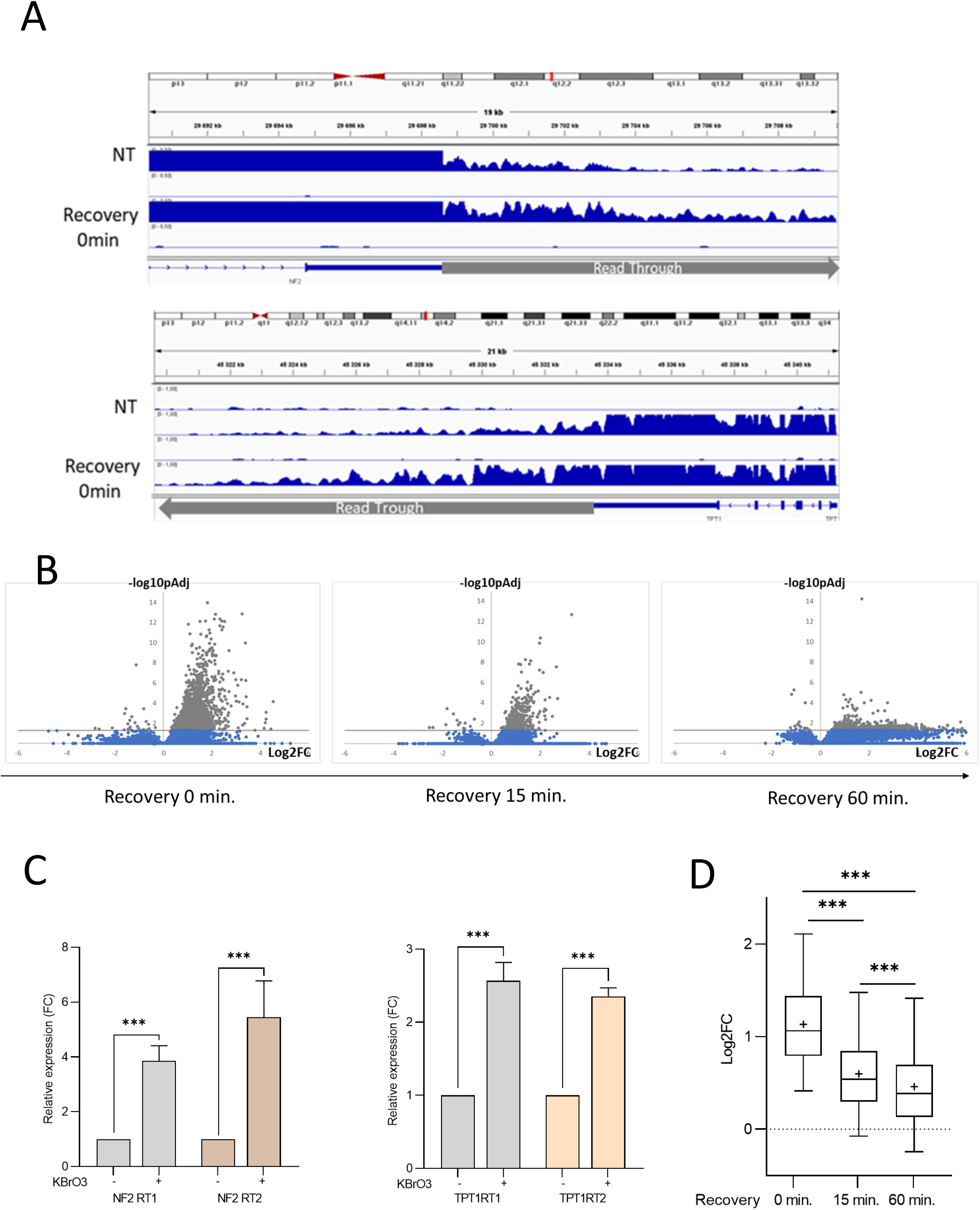
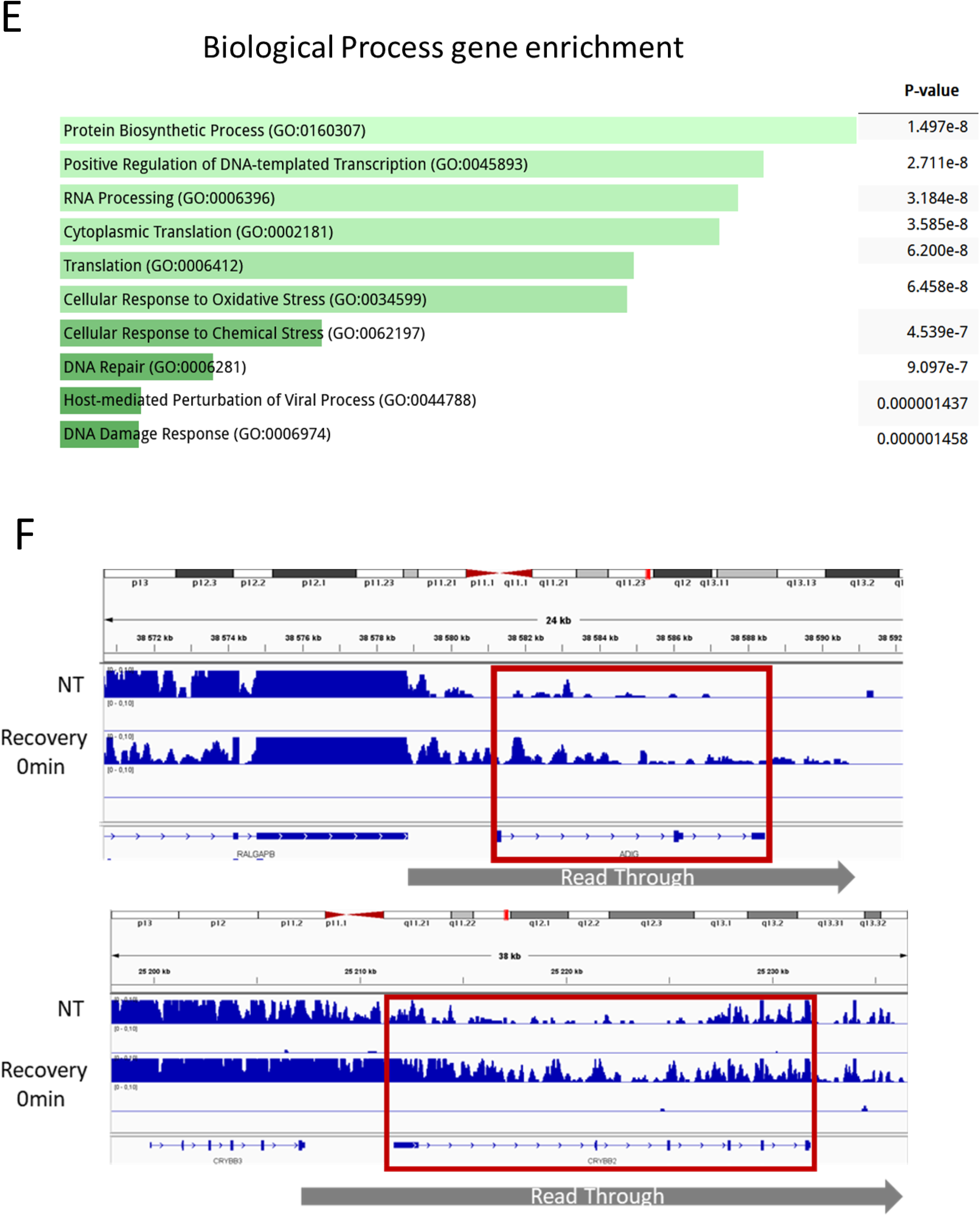
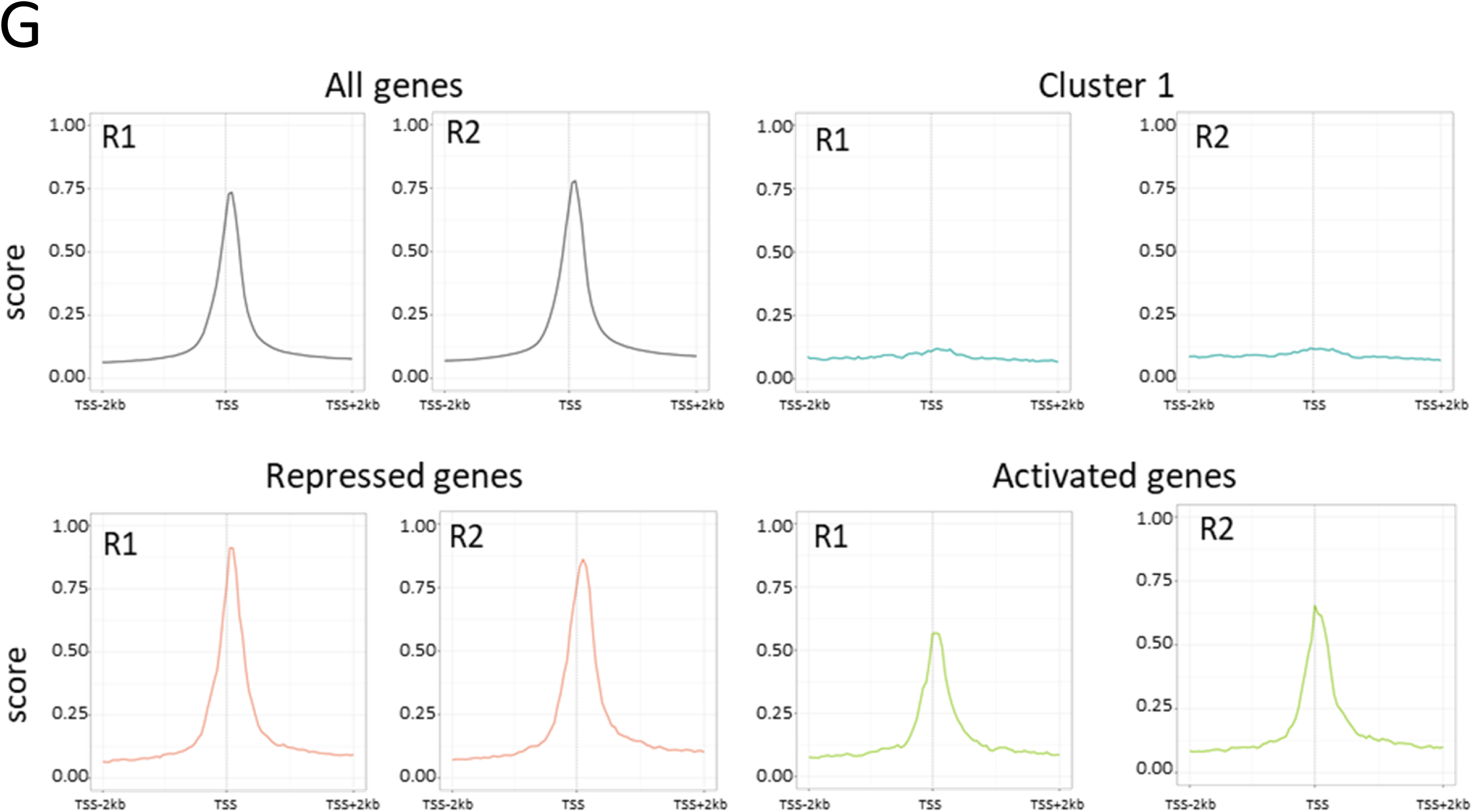
Oxidative stress leads to the activation of read-through RNAs. **A**. Examples of IGV profiles from RNAseq illustrating the activation of readthrough RNAs at early time after KBrO3 treatment. **B.** Volcano plots represent the results of differential expression analyses on read-through RNAs comparing all KBrO3-treated samples to untreated control. **C**. OGG1-FKBP-HA cells were treated or not with KBrO3 for 30 minutes. Total RNA was extracted and analysed by RT-qPCR for the Read-through of NF2 et TPT1 genes as well as the house-keeping RPLP0 pre-mRNAs expression. RT1 and RT2 correspond to the amplification of the proximal and distal read-trough regions, respectively, starting from the TSS. The positions are shown in panel A (red lines). Results were normalized to RPLP0 expression and standardized relative to 1 in untreated cells. The mean and SDOM from three independent experiments are shown. The stars indicate a p-value (<0.05 for *, <0.01 for ** and <0.001 for ***) (Mann-Whitney test). **D**. Box plots with median, 25 and 75% quantiles showing the evolution of genes presenting read-through during recovery following KBrO3 withdrawal. The stars indicate a p-value (<0.05 for *, <0.01 for ** and <0.001 for ***) (Wilcoxon paired test). **E.** GO enrichment analysis for Biological Process terms was performed on genes exhibiting read-through induction in response to KBrO₃ treatment (0 min) using Enrichr. The length of each bar represents the statistical significance of the corresponding gene set or term, while the color intensity (from light to vivid) indicates higher significance. **F**. Examples of IGV profiles illustrating genes from cluster 1 covered by an activated read-through RNA originating from an upstream gene. **G**. OGG1-FKBP-HA cells were subjected to a ChIP Seq using antibodies against Ser5-P RNA polymerase II. Median metaprofiles from TSS-2kb to TSS+2kb are shown for all expressed genes and genes from the three clusters. Two independent replicates (R1 and R2) were analyzed.

### OGG1 is important for genome expression changes upon oxidative stress

We next investigated the consequence of OGG1 depletion on the dynamic transcriptional response to oxidative stress by comparing cells expressing OGG1 (untreated with dTAG) to those depleted of OGG1 (treated with dTAG). We found that genes that are rapidly activated or rapidly repressed in response to oxidative stress were less affected when OGG1 is degraded (Figure 5A and 5B, respectively), indicating that their transcriptional changes are, at least in part, OGG1-dependent. The effect of OGG1 depletion was particularly pronounced for genes rapidly repressed upon oxidative stress induction, with the median Log2 fold change increasing from approximately -1 to -0.5 in the repressed gene population (Figure 5B). In contrast, genes activated only at 1 hour following oxidative stress treatment removal (late activated genes) were not affected by OGG1 depletion (Figure 5C). Thus, these data indicate that OGG1 is required, at least in part, for the rapid regulation of gene expression following oxidative stress induction. Importantly, since OGG1 specifically processes oxidized guanines in DNA these results also suggest that the observed genome-wide expression changes are a consequence of DNA oxidation.

**Figure 5.**
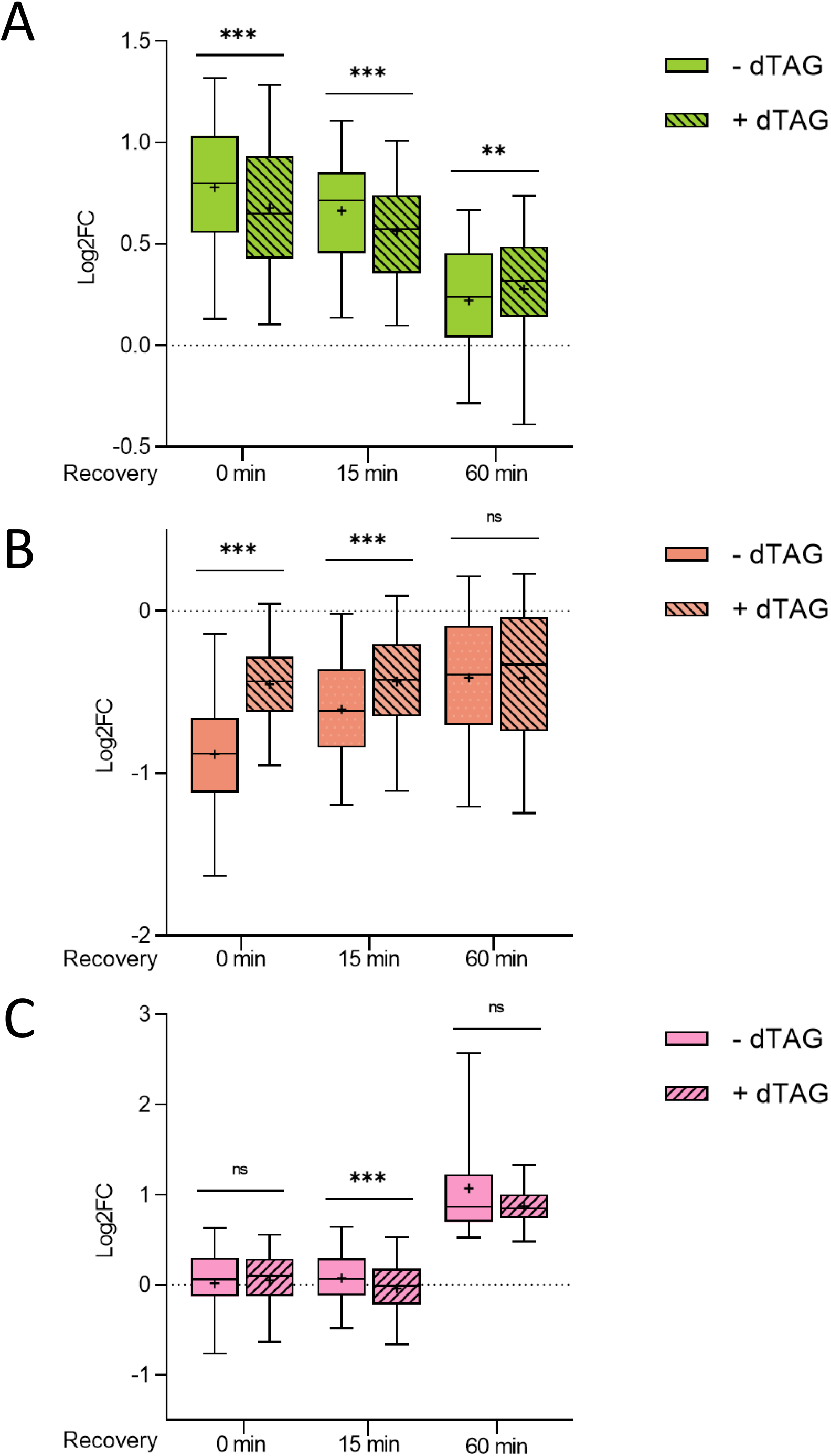
OGG1 influences transcription in response to oxidative stress. Box plots with median, 25 and 75% quantiles showing the evolution of activated genes, repressed genes and late activated genes from Figure 3 clustering during recovery following KBrO3 withdrawal in the presence (-dTAG) and upon induced degradation (+ dTAG) of OGG1. **A**. activated genes. **B**. repressed genes. **C.** late activated genes. The stars indicate a p-value (<0.05 for *, <0.01 for ** and <0.001 for ***) (Wilcoxon paired test)

### Oxidative DNA damage affects RNA pol II recruitment to some promoters

Gene transcription levels can be regulated either through RNA polymerase II (Pol II) recruitment to chromatin or its release from promoter-proximal pausing. To identify which of these steps is affected by oxidative stress, we performed in duplicate ChIP-seq analyses for total RNA Pol II and RNA Pol II phosphorylated on Ser5 (S5-P), which primes Pol II for transcription initiation using the same experimental setting as for RNA-Seq. We next computed total RNA Pol II or S5-P RNA Pol II occupancy at transcription start sites (TSSs), and performed differential gene expression (DGE) analyses of this parameter comparing all samples to the untreated control to assess the effect of oxidative DNA damage and OGG1 depletion. We also calculated the escape index (pol II occupancy over the gene body divided by the sum of pol II occupancy across the TSS and gene body), which reflects promoter proximal pausing release, for untreated cells and cells harvested at 0 minute after KBrO₃ treatment.

We then analyzed these two parameters for the rapidly repressed and activated genes identified in our RNA-seq data. We found that repressed genes exhibited a significant decrease in both total RNA Pol II and S5-P RNA Pol II occupancy at TSSs (Figure 6A), whereas conversely, this occupancy increased for activated genes (Figure 6A). As described previously ^10^, we observed a slight increase in the escape index for all genes compared to the untreated condition (Figure 6B and S4A). However, this increase was largely similar for either activated or repressed genes. Thus, altogether, these data indicate that the rapid transcriptional repression and activation in response to oxidative stress are primarily driven by changes in RNA Pol II recruitment to gene promoters.

**Figure 6.** RNA PolII recruitment under oxidative stress. **A** OGG1-FKBP-HA cells treated with KBrO3 and dTAG as described in Figure 2A were subjected to a ChIPseq analysis using antibodies against total RNA polymerase II (Total Pol II) or S5-P RNA polymerase II (S5-P Pol II). Total Pol II or S5-P Pol II occupancy at TSS were computed for each gene in each condition, and subjected to a differential analysis compared to untreated control. Box plots show the median, 25 and 75% quantiles of the Log2FC of total Pol II or S5-P Pol II occupancy at the 0 minute recovery time point for all genes as well as for rapidly repressed and rapidly activated genes identified in the RNA-Seq. The stars indicate a p-value (<0.05 for *, <0.01 for ** and <0.001 for ***) (Wilcoxon unpaired test). **B**. From the ChIP-seq experiments described in A, the escape index (amount of RNA pol II in the gene body divided by the total amount of RNA pol II at the TSS and the gene body) reflecting promoter proximal pausing release was computed for each gene. Box plots represent the median, 25 and 75% quantiles of this escape index for all genes as well as for rapidly repressed and rapidly activated genes identified in the RNA-Seq, for untreated cells (NT) or cells recovered immediately after KBrO3 treatment (KBrO3 0 min). The red stars indicate a p-value (<0.05 for *, <0.01 for ** and <0.001 for ***) (Wilcoxon unpaired test) between cluster. The black stars indicate a p-value (<0.05 for *, <0.01 for ** and <0.001 for ***) (Wilcoxon paired test) depending of treatment. An independent replicate is shown in Figure S3A. **C**. The differential analysis of RNA pol II occupancy described in A was subjected to unsupervised clustering (see Figure S3C) leading to the identification of genes with increased RNA pol II occupancy (left, in purple) or decreased RNA-Pol II occupancy (right, in brown). Box plots show the median, 25 and 75% quantiles of the Log2FC of RNA pol II occupancy at TSS in the absence (clear boxes) or presence of dTAG (lined boxes) during recovery after KBrO3 treatment for these two categories of genes. The stars indicate a p-value (<0.05 for *, <0.01 for ** and <0.001 for ***) (Wilcoxon paired test). **D**. Same as in C, except that S5-P RNA pol II ChIPseq data were used. Unsupervised clustering is shown in Figure S3D. The stars indicate a p-value (<0.05 for *, <0.01 for ** and <0.001 for ***) (Wilcoxon paired test).

Strikingly, when analyzing the DGE at the 0 minute time point, many more genes exhibited significant changes in RNA Pol II recruitment to TSSs compared to the number of genes with detectable changes in intron expression in RNA-seq (compare the volcano plot in Figure S4B with that in Figure 2B). This suggests that oxidative stress-induced changes in RNA Pol II recruitment do not always translate into detectable changes in intron expression, likely because they are transient and partially masked by the half-life of introns. We therefore recovered all genes for which RNA Pol II occupancy at TSSs was significantly affected with a Log2FC >1 or <-1 in at least one condition and performed a clustering analysis as described above. We identified two main clusters of genes: one in which RNA Pol II occupancy rapidly increases and another in which it rapidly decreases (Figure S4C). We did not identify genes for which RNA Pol II occupancy increases at 1 hour, which could correspond to the late-activated genes identified in RNA-seq, likely because the magnitude of changes in RNA Pol II occupancy was insufficient for detection. When analyzing the evolution of RNA Pol II occupancy over time in these two clusters, we observed a pattern strikingly similar to that in total RNA-seq, with a rapid return to baseline levels for both clusters (Figure 6C). Analyzing the RNA Pol II occupancy after dTAG treatment, we observed that the decrease in RNA Pol II occupancy upon oxidative stress is OGG1-dependent, consistent with our RNA-seq observations (Figure 6C), whereas no significant effect of OGG1 depletion on the increase in RNA pol II occupancy could be seen. A similar analysis of S5-P Pol II ChIP-seq data yielded very comparable results (see Figure S4D for clustering and 6D for analysis of clusters).

Thus, these data indicate that oxidative stress affects RNA Pol II occupancy at many promoters, either positively or negatively, resulting in changes in transcription levels. Importantly, the decrease in RNA Pol II occupancy at gene promoters is, at least in part, OGG1-dependent and thus probably linked to oxidative DNA damage, confirming our RNA-seq results.

### Setting up a system allowing induction of local oxidative DNA damage

The results from the above-described experiments indicate that oxidative DNA damage leads to genome-wide expression changes, including the rapid activation or repression of specific genes. We next sought to determine whether these changes could be due to local oxidative DNA damage. Indeed, previous studies using transient transfection of *in vitro* oxidized reporter DNA sequences have shown that DNA oxidation can directly affect transcription ^24^. To address this question for endogenous genes, we adapted a recently described optogenetic cellular tool to target oxidative DNA damage to specific genomic loci ^25^ ; ^26^. For this purpose, we used the fluorogen activating protein (FAP), which generates reactive oxygen species (ROS) in the presence of diiodinized malachite green (MG2i) upon light activation ^27,28^ . Notably, this FAP has already been used in a fusion protein with TRF1 to generate oxidative DNA damage specifically at telomeres ^25^ ^26^. We fused FAP to the catalytically inactive Cas9 (dCas9) protein to target it to selected genomic regions using specific guide RNAs (gRNAs). We generated a stable U2OS cell line (FAP-dCas9 cells) expressing this fusion protein (see Figure 7A for a schematic description). To validate this tool, we transfected FAP-dCas9 cells with gRNAs targeting telomeric repeats. This resulted in the recruitment of the FAP-dCas9 fusion protein to telomeres labelled by an anti-TRF1 antibody (Figure 7B). We next added MG2i, illuminated cells and followed by live microscopy the dynamics of OGG1-GFP and FAP-dCas9 (to follow telomeres). Although illumination resulted in rapid FAP-MG2i signal bleaching, telomeres positions are still readily detectable 30 s after illumination (Figure 7C). Most importantly, upon the addition of MG2i, we observed the light-induced recruitment of OGG1-GFP to FAP-dCas9 foci (Figure 7C). Quantification of OGG1-GFP images showed rapid and specific recruitment to telomeres (Figure 7D and E). Since OGG1 specifically recognizes oxidized bases, these data demonstrate that we successfully generated oxidative DNA damage at telomeres. Therefore, this result indicates that FAP-dCas9 cells enable the induction of oxidative DNA damage that induce OGG1 recruitment at selected genomic sequences in a spatio-temporally controlled manner.

**Figure 7.**
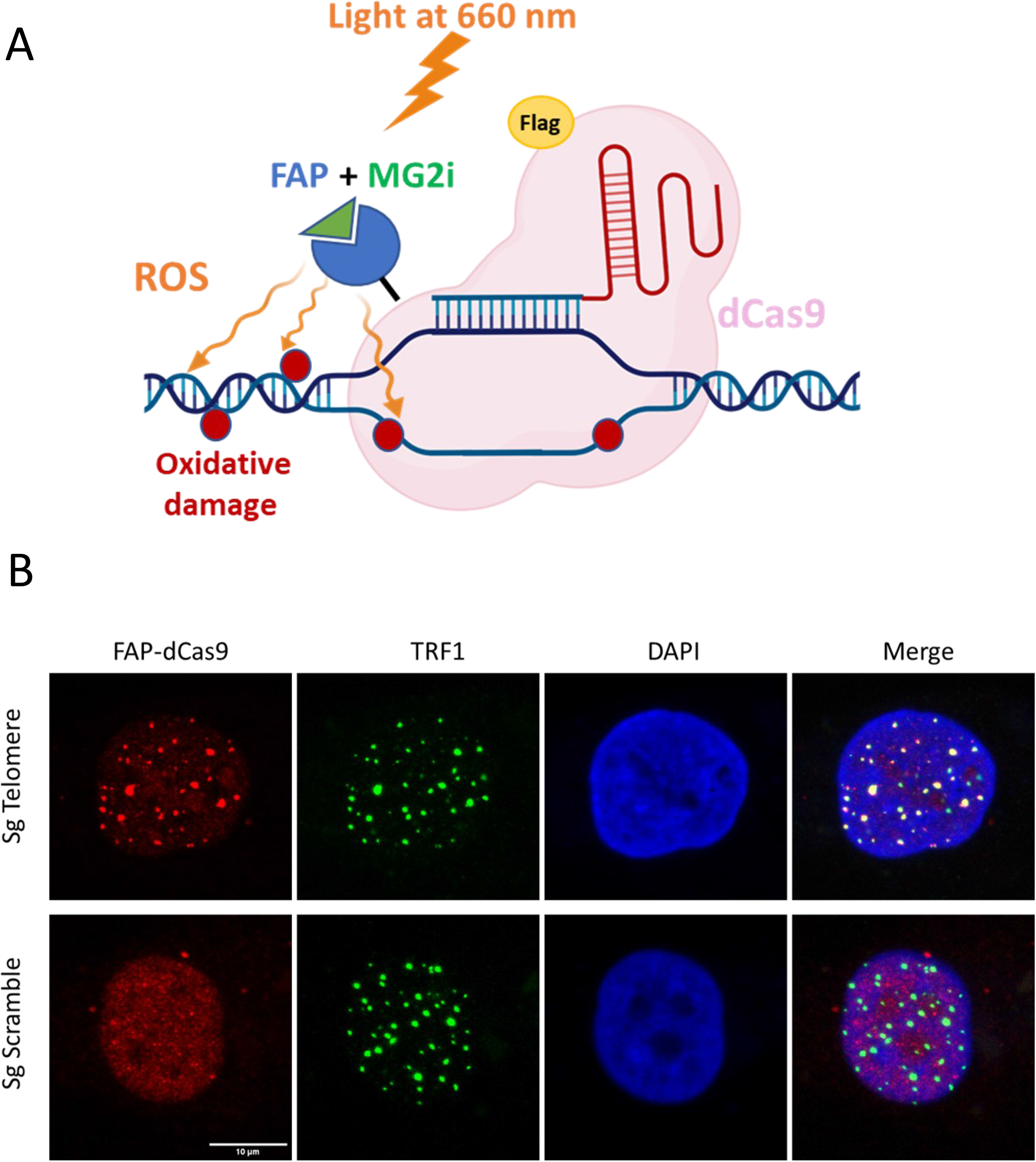
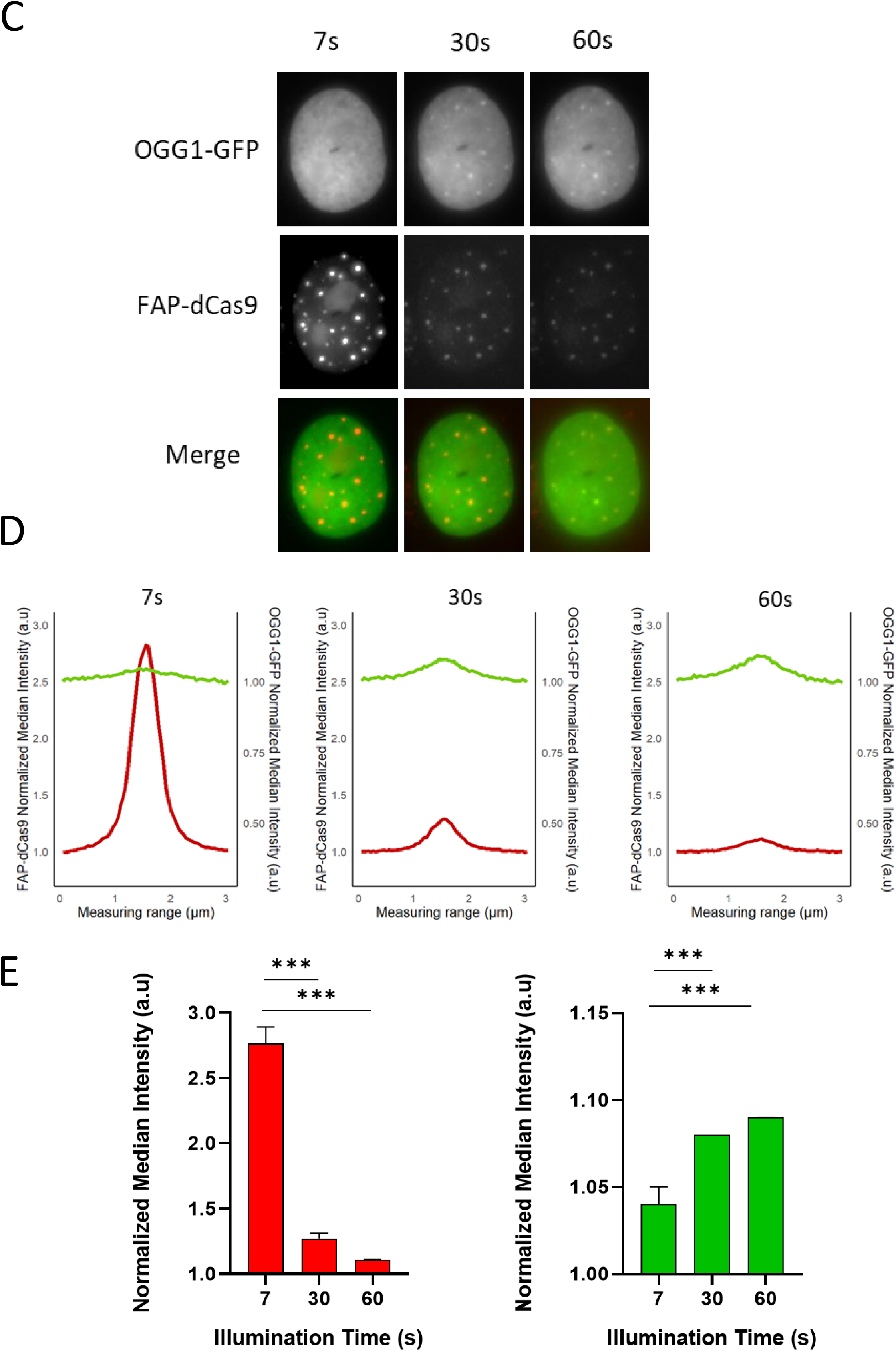
Targeted generation of ROS on the genome. **A**. Schematic representation of the optogenetic system allowing the targeted production of ROS. **B**. Representative images showing localization of FAP-dCas9 at telomeres. FAP-dCas9 cells transfected with a sgRNA targeting telomeric sequence or a control sgRNA (scramble). Telomeres localization was revealed by TRF1 immunostaining and FAP-dCas9 by Flag immunostaining. (scale bar: 10µm). **C**. FAP-dCas9 cells stably expressing OGG1-GFP were transfected with sgRNAs targeting telomeric sequences, incubated with MG2i for 30 min then exposed to red light at 660nm. Images showing MG2i fluorescence (representing FAP-dCas9 localization) and OGG1-GFP were acquired during the first illumination, 30 and 60 seconds following illumination. A representative cell is shown. **D.** Kinetic on live cells imaging showing the recruitment of OGG1-GFP at FAP-dCas9 foci after ROS generation. The red curve is the median signal obtained from 180 foci of MG2i -FAP-dCAs9 and the green curve is the median intensity of OGG1-GFP foci for the same foci. **E**. Quantification of the median intensity for MG2i-FAP-dCas9 and OGG1-GFP from live cell experiments. Represented are the normalized median intensity. The stars indicate a p-value (<0.001 for ***) (Wilcoxon paired test).

### Targeting oxidative DNA damage to specific regions induced local genome expression changes

We next aimed to target oxidative DNA damage to promoters that we identified in this study as rapidly affected by oxidative DNA damage treatment to assess whether their transcriptional changes are due to local DNA oxidation. To this end, we designed guide RNAs (gRNAs) targeting the *HAPLN1* and *CAV1* promoters, which are repressed upon KBrO₃ treatment, and the *SLC41* and *NCKAP5L* promoters, which are activated by KBrO₃ treatment (see Figure 2).We next transfected these gRNAs into the FAP-dCas9 cell line. Chromatin immunoprecipitation (ChIP) experiments with Cas9 antibodies confirmed the targeting of the FAP-dCas9 fusion protein to their respective target promoters upon transfection of the specific gRNAs (Figure S5). We then treated cells with or without MG2i, illuminated them, and performed RT-qPCR on introns of the genes. Strikingly, we found that targeting the *HAPLN1* and *CAV1* promoters resulted in a MG2i-dependent decrease in *HAPLN1* and *CAV1* pre-mRNA expression in response to illumination (Figure 8A and B), thereby mimicking the effect observed upon KBrO₃ treatment. In contrast, no MG2i-induced activation of transcription was observed for *SLC41A1* and *NCKAP5L* (Figure 8C and D). This result could suggest that the activation of these promoters is not due to local oxidative DNA damage. However, we cannot rule out the possibility that we did not achieve enough well-localized oxidative DNA damage at these two promoters to detect a direct effect of oxidation. Nevertheless, at least for the *HAPLN1* and *CAV1* repressed promoters, the effect of oxidative stress is due to direct oxidative DNA damage around their promoters. These data demonstrate, for the first time, that oxidative stress can induce rapid genome-wide expression changes at endogenous promoters due to local DNA oxidation.

**Figure 8.**
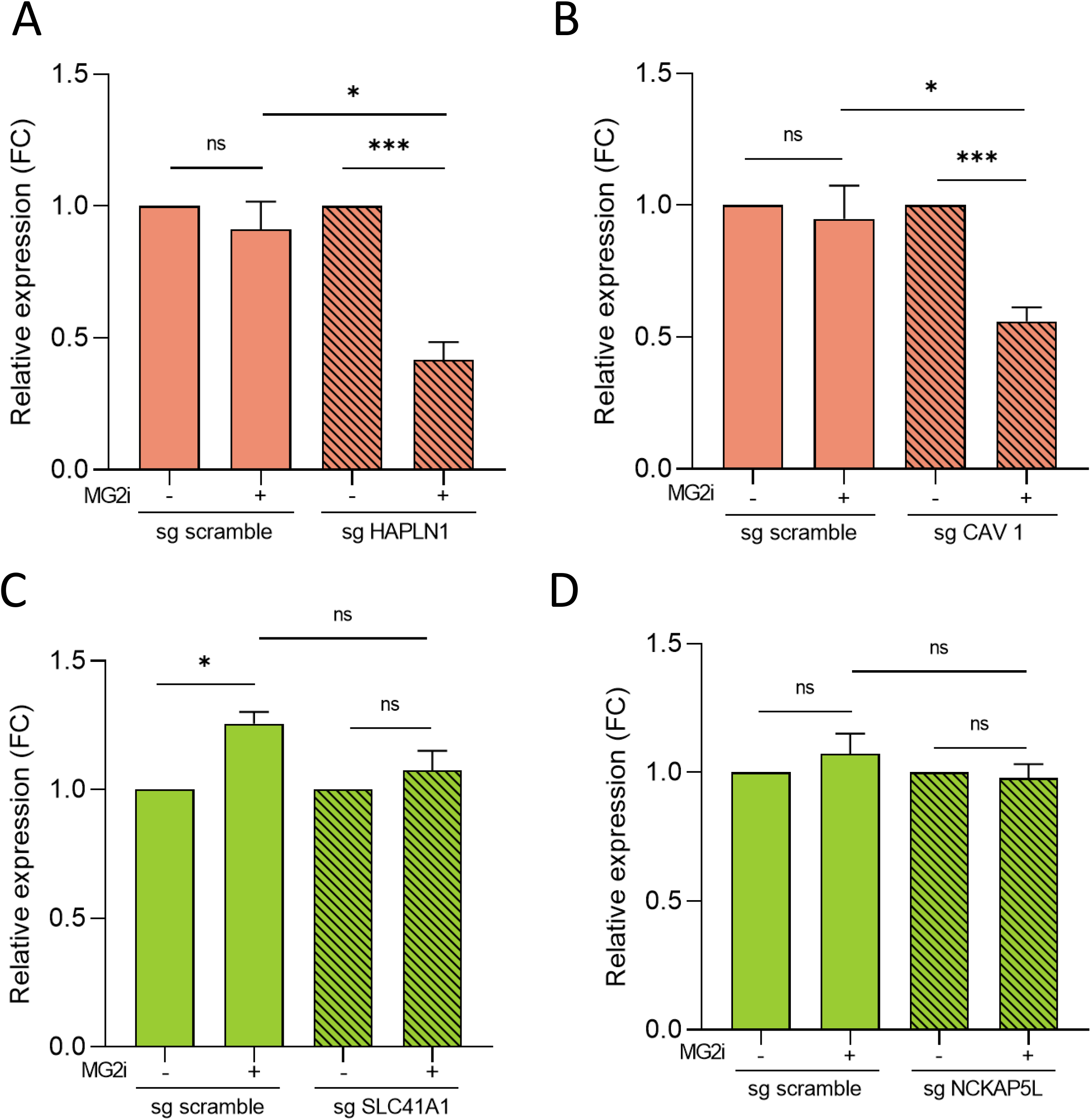
Local induction of oxidative DNA damage represses transcription of the CAV1 and HAPLN1 gene. **A**. FAP-dCas9 cells were transfected by the indicated sgRNAs. 24 hours following transfection, MG2i was added or not, as indicated and cells were illuminated at 660nm to induce ROS. Total RNAs were extracted 15 minutes later and analysed by RT-qPCR for HAPLN1 as well as the house-keeping RPLP0 premRNAs expression. Results were normalized to RPLP0 expression and standardized relative to 1 in the absence of MG2i. The mean and SDOM from 3 independent experiments are shown. Statistical analysis was done with student t-test. **B.** Same as in A, except that sgRNAs targeting CAV1 were used. CAV1 premRNA expression was analysed as described in A. The mean and SDOM from 3 independent experiments are shown. **C.** Same as in A, except that sgRNAs targeting SLC41A1 were used. SLC41A1 premRNA expression was analysed as described in A. The mean and SDOM from 3 independent experiments are shown. **D.** Same as in A, except that sgRNAs targeting NCKAP5L were used. NCKAP5L premRNA expression was analysed as described in A. The mean and SDOM from 3 independent experiments are shown. The stars indicate a p-value (<0.05 for *, <0.01 for ** and <0.001 for ***) (Wilcoxon paired test).

## Discussion

Here, we evidenced four features of transcriptional response to oxidative stress: first, a rapid activation or repression of many genes; These changes are transcriptional, and gene repression is partially dependent on OGG1 and mediated by direct local oxidation of promoters. Second, a recovery of transcription of these genes about one hour after stress removal; third, a delayed activation of some genes which probably reflects the setting up of the response to oxidative stress; fourth, a rapid activation of read-through RNA expression.

By ChIPseq against RNA pol II, we found that the effects of oxidative stress on promoter activity are mostly due to changes in RNA pol II recruitment to the TSS of some genes. Although we found that oxidative DNA damage weakly induces release of RNA pol II from promoter proximal pausing, consistent with what has been shown following H_2_O_2_ treatment ^9,10^, this release is not different for genes which are activated and repressed. We can speculate that depending on the promoter context, DNA oxidation or its management can favor or disfavor the assembly of the pre-initiation complex.

We also found that oxidative stress rapidly induced the expression of thousands of read-through RNAs which are produced downstream of genes due to defects in transcription termination. Although activation of read-through RNAs expression by oxidative stress was already shown ^11^, these studies were performed at much longer time following oxidative stress. Our data suggest that oxidative stress may rapidly affect transcriptional termination. We observe a weak effect of OGG1 depletion on this feature, which could indicate a contribution of DNA oxidation in this process. How DNA oxidation can affect transcriptional termination remains to be determined. Importantly, read-through RNAs have been proposed to be important for maintaining the stiffness of the nucleus upon stress ^29^. This general and rapid activation of read-through RNAs may be functionally important. Strikingly, we found that genes from which these read-through RNAs originate are enriched in genes from the response to oxidative DNA damage. This suggests that upon oxidative stress, transcription termination of these genes may be altered, leading to the production of read-through RNAs. Whether some of these read-through RNAs could have specific functions, as other long non-coding RNAs, in the response to oxidative stress is an attractive hypothesis.

Importantly, we investigated the function of OGG1 in gene expression changes upon DNA damage. Until now, the function of OGG1 in transcription has been mainly studied using OGG1 negative cells ^9^, in which one cannot rule out the possibility of cell adaptation to OGG1 depletion with the activation of alternate pathways for 8-oxoG management as already suggested ^30,31^. Results obtained with OGG1 inhibitors, as used in ^32^, have also to be interpreted with great care since it was recently shown that these molecules presented off target effects, questioning their specificity ^16^. Thanks to an inducible-degron based system used on endogenous OGG1, we were able to investigate here the consequences of a fast acute OGG1 depletion, avoiding cell adaptation to OGG1 depletion observed in KO cells. This allowed us to demonstrate that OGG1 is important for the rapid repression of specific genes observed upon oxidative DNA damage management. These results indicate that repression of these genes is a consequence of DNA oxidation. However, repression is reduced in the absence of OGG1, indicating that DNA oxidation *per se* is not responsible for this repression. This is consistent with the ability of RNA Pol II to bypass some oxidative DNA damage ^8^. Rather, the recognition and processing of oxidized bases initiated by glycosylases such as OGG1 interferes with the normal functioning of promoters. For example, the presence of OGG1 and other BER proteins may sterically hinder the normal function of these promoters. We can speculate that in the absence of OGG1, recognition or processing of other oxidized bases initiated by the other glycosylases is responsible for the residual transcriptional repression.

One striking behavior observed following oxidative stress removal is the rapid recovery of normal transcription levels, since the effect of oxidative stress is nearly abolished one hour following removal and already strongly decreased at 15 minutes time point. This suggests that cells developed rapid and potent mechanisms which reset promoters following oxidative DNA damage repair. These mechanisms are probably essential for cells in order to avoid long term consequences of oxidative DNA damage on gene expression. 8-oxoG mapping following KBrO3 removal in U2OS cells has already shown that the repair of oxidative DNA damage is particularly efficient at active promoters, with a decrease of 8-oxoG levels already observed 1 hour following KBrO_3_ removal ^33^ revealing a coupling between repair of oxidative DNA damage and transcription. This kinetic is strikingly similar to the resetting of gene transcription we show here. We can speculate that such a rapid resetting of gene transcription also involves a coupling between repair mechanism and transcription restart. This is in agreement with recent data showing the importance of OGG1 in transcription restart ^7^.

Finally, a question that remained unanswered in the field is whether gene expression changes upon oxidative DNA damage are due to direct DNA oxidation at gene promoters. Most importantly, to answer this question, we developed a unique system allowing to induce oxidative DNA damage at targeted specific sequences. This can allow to study the consequences of local oxidative DNA damage independently of checkpoints induction or activation of intracellular signaling pathways due to massive and global oxidative stress when chemical treatment is used (such as H2O2, KBrO_3_, menadione…). We validated this tool by showing that it allows the generation of oxidative DNA damage at unique sites in the genome, since we can recapitulate on two genes the effect of global oxidative stress. This system represents a unique and versatile tool that will be very valuable to understand the relationship between oxidative DNA damage management and chromatin landscape. We illustrate here the major interest of this system that allowed us to demonstrate for the first time to our knowledge that local oxidative DNA damage can affect the expression of endogenous genes.

## Materials and Methods

### Tagging of endogenous OGG1 protein for inducible degradation

PX458_sgOGG1 plasmid: guide RNA sequence (AAGGGTTCCAAAGGGCCGGA) targeting the coding sequence upstream the stop codon of the OGG1 gene (variant 1a, NM_002542) has been inserted between the BbsI restriction sites of the plasmid pSpCas9(BB)-2A-GFP (PX458), a gift from Feng Zhang (Addgene plasmid # 48138; http://n2t.net/addgene:48138; RRID: Addgene_48138)

HDR Donor plasmid: FKBP^F36V^-2xHA-P2A-BSD fragment has been amplified from the plasmid pCRIS-PITChv2-dTAG-BSD (BRD4), a gift from James Bradner & Behnam Nabet (Addgene plasmid # 91795; http://n2t.net/addgene:91795; RRID: Addgene_91795). Left (214 nt) and right (329 nt) homology sequences on both sides of the OGG1 gene stop codon have been amplified from U2OS cells genomic DNA. All PCR primers have been designed with overlapping ends necessary for Gibson assembly (NEB, E5510) in the pBSk+ plasmid at the EcoRV restriction site.

For U2OS-OGG1-FKBP-HA cell line generation, U2OS cells from ATCC have been co-transfected with PX458_sgOGG1 and the HDR donor plasmid using JetPrime transfection reagent respecting the standard plasmid DNA transfection protocol from Polyplus. 48h after transfection, cells have been diluted and treated with Blasticidin antibiotic (Invivogen) at 8µg/ml. Resistant clones have been selected for the correct insertion of FKBP-v-2xHA-P2A-BSD in both alleles via out/out PCR and sequencing.

### Establishment of U20S cell line expressing FAP-hdCas9

The lentiviral vector pTRIP_3XFlag_dL5_2X4GS_hdCas9_P2A_BSD was obtained by cloning. Briefly, 3XFlag-SV40 NLS -dCas9-Nucleoplasmin NLS sequence has been amplified from the plasmid BIC-Gag-dCas9, a gift from Aline Marnef (CBI, Toulouse, France) which introduced D10A and H840A mutations in the Cas9 sequence of the plasmid BIC-Gag-Cas9 (Addgene plasmid # 119942; http://n2t.net/addgene:119942 ; RRID:Addgene_119942). The DNA fragment containing dL5-2X4GS come from the plasmid pcDNA3.1-NLS-myc-dL5-2xG4S-mCer3, a gift from Marcel Bruchez (Addgene plasmid # 73205 ; http://n2t.net/addgene:73205 ; RRID:Addgene_73205) and P2A-BSD DNA come from the plasmid pCRIS-PITChv2-dTAG-BSD (BRD4), a gift from James Bradner & Behnam Nabet (Addgene plasmid # 91795 ; http://n2t.net/addgene:91795 ; RRID:Addgene_91795)

All DNA fragments have been amplified by PCR using primers with overlapping ends allowing Gibson assembly (NEB, E5510) in the pTRIP-eF1α-GFP lentiviral vector backbone, a gift from Loïc Vandenbergue (CRCT, Toulouse, France) opened with BAMH1 and XhoI restriction enzyme in order to remove GFP.

### Cell treatments and illumination

To induce the degradation of the endogenous OGG1 protein, the dTAGV-1 molecule (HY-145514, MedChemExpress) diluted in DMSO is added in the culture medium at 500 nM for the indicated time. Same volume of DMSO is used for control conditions.

To generate acute oxidative stress, KBrO3 (Sigma) was added in DMEM GlutaMAX medium (Gibco) supplemented with 10% SVF and 1 mM sodium pyruvate (Gibco) at 40mM for 30 min. To generate targeted singlet oxygen, cells were incubated in FluoroBrite DMEM medium (Gibco) supplemented with 10% SVF, 1X GlutaMAX (Gibco) and 1 mM sodium pyruvate (Gibco) containing 100 nM of the stable fluorogen MG2i (generous gift from Brigitte Schmidt (Mellon University, Pittsburg) for 30 min at 37 ° C, 5% CO_2_ and 5% O_2_ before exposition to 660 nm LED light (Source CoolLED pE-100, Optoprim) at 100 mW/cm2 for 15 min (unless indicated otherwise).

### Cell Culture and SgRNA transfection

U2OS cells were grown in DMEM containing Glutamax (Gibco) supplemented with sodium pyruvate and 10% fetal bovine serum under 5% CO_2_ and 5% O_2_

Cells are transfected with 3nM of the different SgRNA or 25nM for sgRNA targeting telomeres using DharmaFECT 4 reagent (Horizon discovery) for 15h.

### Western blot

Cell pellets were resuspended in lysis buffer (2% SDS, 10% glycerol, 60 mM Tris-HCl pH 6.8, supplemented with protease and phosphatase inhibitors) and incubated on ice for 30 minutes. The suspension was then sonicated for 1 minute (30% amplitude, 1-second ON/1-second OFF pulses). Protein concentration was quantified using the Qubit Protein BR Assay. Thirty micrograms of protein were resolved by SDS-PAGE on a 4–12% Bis-Tris polyacrylamide gel (NuPAGE 4–12%, Invitrogen) and transferred onto a 0.2 µm nitrocellulose membrane.

Membranes were blocked with 5% skim milk in TBS buffer containing 0.01% Tween-20 and incubated overnight at 4°C with primary antibodies diluted in the blocking solution. After washing with TBS-T (0.02% Tween-20), membranes were incubated with HRP-conjugated secondary antibodies in blocking solution for 1 hour at room temperature, followed by additional washes in TBS-T (0.02% Tween-20). HRP signal detection was performed using ECL Western Blotting Substrates (Clarity Max, Bio-Rad), and visualization was carried out with the ChemiDoc Imaging System (Bio-Rad).

### EU incorporation, staining and acquisition

A total of 10,000 cells were seeded in a 96-well glass-bottom plate in complete medium. After 24 hours, cells were treated with either 40 mM KBrO₃ or 1 mM H₂O₂ (freshly prepared in complete medium) for 30 minutes at 37°C, 5% CO₂, and 5% O₂. 5-ethynyl uridine (EU) at 750 µM was added during the treatment.

EU incorporation was detected using the Click-iT RNA Alexa Fluor 488 Imaging Kit (Invitrogen). Briefly, cells were fixed with 4% paraformaldehyde (PFA) in PBS for 20 minutes, washed three times with PBS, and permeabilized with 0.5% Triton X-100 in PBS for 15 minutes. After two additional PBS washes, cells were incubated with the 1× reaction mix (prepared according to the manufacturer’s instructions) for 1 hour. Following three PBS washes, nuclei were counterstained with DAPI (10 µg/mL) for 10 minutes.

Fluorescence images were acquired using a Zeiss Cell Discoverer 7 high-content microscope and analysed with Arivis software.

### Immunofluorescence

Cells were seeded on coverslips, transfected or treated as indicated. Cells were washed with PBS and fixed at room temperature with 4% formaldehyde (PFA) for 10 min. Fixed cells were washed with PBS, permeabilized with PBS-Triton 0.5% during 7 min before blocking with 3% BSA in PBS. Cells were incubated overnight at 4 °C with indicated primary antibodies. The next day, cells were washed 3 times with PBS, incubating with Alexa Fluor coupled secondary antibodies (Thermo Fisher) 1h at room temperature and washing again 3 times with PBS. Nuclei are counterstained in a solution of PBS with 1 µg/ml of DAPI. After 2 washes in PBS and 1 in water, coverslips were mounted in vectashield antifade mounting medium. Images were acquired using a Leica SP8 confocal microscope with a 40X objective lens.

### OGG1 protein recruitment Imaging assay

Recruitment assays for DNA repair proteins were performed using live-cell imaging to monitor the dynamics of protein recruitment in real time. In both approaches, FAP-dCas9 was specifically activated using 660-nm illumination in presence of MG2i, as detailed below.

Cells were seeded on coverslips, transfected or treated as indicated. To follow the kinetics of OGG1-GFP protein recruitment, cells were imaged in FluoroBrite medium (phenol-red free) supplemented with 10% FBS, sodium pyruvate, HEPES, GlutaMAX, and penicillin–streptomycin, and maintained at 37°C on a heated microscope stage. In this setup, FAP-dCas9 was activated directly under the microscope, using the Cy5 excitation filter set (excitation 628 ± 40 nm, emission 692 ± 40 nm), which corresponds to the activation wavelength of FAP-MG2i complex. Activation was performed during image acquisition. The first image was acquired during the 7 seconds of acquisition time (8 z stack, 900ms acquisition time each). After activation, time-lapse imaging was performed to monitor real-time recruitment of OGG1-GFP protein.

### Total RNA extraction

Cells were directly lysed in the culture plate with 0.1 mL/cm^2^ of TRIzol (TRI Reagent Sigma) during 5 min, and the lysate was harvested before adding 0.1 mL of chloroform/mL of TRIzol. After centrifugation, the upper phase was collected and RNA is precipitated in propan-2-ol, and washed in 75% Ethanol. The MasterPure RNA Purification Kit (Epicentre) was used for the DNase digestion and subsequent precipitation. After total nucleic acids recovery, DNA was removed with a cocktail of DNase I and Baseline Zero DNase supplemented with Riboguard RNase inhibitor for 45 min at 37°C.

### Reverse transcription

A 500-ng amount of total RNA was reverse transcribed with SuperScript III reverse transcriptase (Invitrogen) according to the manufacturer’s instructions. A condition without SuperScript III was included for each sample and analyzed by qPCR using GAPDH intron 8 primers to check for DNA contamination. qPCR was performed in triplicate with TB green Premix *Ex Taq* from TaKaRa and the Bio-Rad CFX thermocycler. RPLP0 premRNA expression was used to normalize RT-qPCR results.

### Nascent RNA labelling and capture

Nascent RNA was labelled and captured using the Click-iT™ Nascent RNA Capture Kit (Invitrogen, Thermo Fisher Scientific, Cat. C10365) according to the manufacturer’s instructions, with minor modifications. U2OS cells were seeded at 5 × 10⁵ cells per 6-cm dish 24 h before labeling. For EU incorporation, cell culture medium was supplemented with 5-ethynyl uridine (EU, Component A) at a final concentration of 0.2 mM (1:500 dilution from a 100 mM stock in DMSO) and cells were incubated at 37 °C in a humidified 5% CO₂ incubator for 30 min (simultaneously the cells were co-treated or not with 40mM KBrO3 for 30 min). Total RNA was extracted following the previously described protocol and quantified by NanoDrop.

For click-chemistry–mediated biotinylation, 2 µg of total RNA in a final volume of 50 µL was incubated with the Click-iT reaction cocktail prepared from kit components as follows: EU buffer 2× (Component B), CuSO₄ solution (Component D), biotin azide (Component C), Click-iT reaction buffer additive 1 (Component E) and RNase-free water were combined according to the manufacturer’s recommendations, and RNA was added last. After a 3-min incubation at room temperature, Click-iT reaction buffer additive 2 (Component F) was added to initiate the copper-catalyzed azide–alkyne cycloaddition. The reaction mixture was incubated for 30 min at room temperature with gentle vortexing to allow efficient conjugation of biotin azide to EU-labeled RNA. Biotinylated RNA was precipitated by adding glycogen (20 mg/mL), 7.5 M ammonium acetate and 100% ethanol, incubating at −80 °C overnight, then centrifuging for 20 min at 13,000 × *g*, 4 °C. Pellets were washed twice with 70% ethanol, air-dried and resuspended in 25 µL RNase-free water.

Capture of nascent RNA was performed using Dynabeads™ MyOne™ Streptavidin T1 magnetic beads (Component H) supplied in the kit. Beads (24 µL per sample) were washed three times in Click-iT reaction wash buffer 2 (Component J) and resuspended in the same buffer. For each reaction, 25 µL of biotinylated RNA was mixed with 95 µL of RNA binding mix containing 2× RNA binding buffer (Component G), RNase inhibitor and RNase-free water, then heated to 69 °C for 5 min to disrupt secondary structures. Beads were added and incubated with the RNA binding mix for 30 min at room temperature with gentle agitation. Bead–RNA complexes were washed five times with Click-iT reaction wash buffer 1 (Component I), followed by five washes with Click-iT reaction wash buffer 2 (Component J), to remove non-specifically bound RNA. Finally, beads were resuspended in 50 µL of Click-iT reaction wash buffer 2 / water.

Reverse transcription was performed directly on bead-bound nascent RNA using SuperScript™ III reverse transcriptase (Invitrogen). Beads were pre-equilibrated at 70 °C for 5 min, and a reverse transcription mix containing random primers, dNTPs, 5× First-Strand buffer, DTT, RNase inhibitor and SuperScript III was added immediately. Reactions (100 µL final volume) were incubated for 1 h at 42 °C, followed by 5 min at 85 °C to inactivate the enzyme. Beads were magnetically separated and the supernatant containing cDNA derived from nascent RNA was collected and analysed by qPCR.

### Total RNAseq

U2OS cells treated or not with KBrO3 and dTAGV1 and followed a recovery kinetic as described. Total RNA extraction was performed followed by library preparation and sequencing. Total RNA integrity was assessed on a Bioanalyzer then a total of 1 µg of intact total RNA per sample was used as input for rRNA depletion with the NEBNext rRNA Depletion Kit v2, following the manufacturer’s instructions. The depleted RNA was purified and fragmented according to the NEBNext Ultra II RNA Library Prep protocol. First- and second-strand cDNA synthesis were performed, followed by end repair, adaptor ligation, and PCR enrichment (number of cycles adjusted according to input). cDNA was purified between steps using magnetic beads as recommended. Final libraries were quantified and quality-controlled by Bioanalyzer, showing intact profiles. Equimolar amounts of each library were pooled prior to sequencing.

### Chromatin Immunoprecipitation

Chromatin immunoprecipitations (ChIPs) were performed on 75 million U2OS cells treated or not with KBrO3 and dTAGv1 and following the same recovery kinetic described on the RNAseq. Cells were first cross-linked for 15 minutes using 1% formaldehyde added directly to the culture medium, followed by the addition of 0.125 M glycine for 5 minutes. After two washes with cold phosphate-buffered saline (PBS), cells were scraped snap-frozen in liquid nitrogen and stored as dry pellets at −80 °C.

Cells were lysed in 3 mL of lysis buffer (5 mM piperazine-N,N′-bis(2-ethanesulfonic acid) (pH 8), 85 mM KCl, 0.5% NP-40) and homogenized with a Dounce homogenizer. After centrifugation, nuclear pellets were resuspended in 1.5 mL of nuclear lysis buffer (25 mM Tris [pH 8.1], 5 mM EDTA, 0.2% SDS) and sonicated 10 times for 10 seconds (power 0.5, 50% amplitude; Branson Sonifier 250) to obtain DNA fragments of approximately 500 bp. DNA concentration was measured using a Nanodrop system, and samples were adjusted to the same chromatin concentration.

Samples were then diluted twofold in dilution buffer (0.01% SDS, 1.1% Triton X-100, 1.2 mM EDTA [pH 8], 17 mM Tris [pH 8.1], 167 mM NaCl) and precleared for 2 hours with 250 µL of 50% pre-blocked protein A/protein G beads (Sigma P-7786 and P-3296, respectively). Bead blocking was performed by incubating them with 0.5 mg/mL ultrapure bovine serum albumin (BSA) for 4 hours at 4 °C. A volume of 100 µL of chromatin was kept aside as input. For each ChIP, 150 µg of precleared samples were incubated overnight with antibody at 4 °C. A no-antibody control sample was processed in parallel. Then, 50 µL of 50% blocked protein A/G beads were added for 2 hours at 4 °C to recover immune complexes.

Beads were washed once with dialysis buffer (2 mM EDTA, 50 mM Tris [pH 8], 0.2% Sarkosyl), three times (a first short 5min wash, followed by two longer 10min wash) with wash buffer (100 mM Tris [pH 8.8], 500 mM LiCl, 1% NP-40, 1% sodium deoxycholate), and twice with TE buffer (10 mM Tris [pH 8], 1 mM EDTA). Bead–chromatin complexes were resuspended in 200 µL TE buffer and incubated for 30 minutes at 37 °C with 10 µg RNase A (Abcam), along with the input DNA. Cross-linking was reversed in the presence of 0.2% SDS at 70 °C overnight with shaking. After a 2-hour treatment with proteinase K (0.2 mg/mL) at 45 °C, immunoprecipitated DNA and input DNA were purified using a phenol/chloroform extraction. All buffers used for ChIP experiments were supplemented with an EDTA-free protease inhibitor cocktail (Roche) and filtered through a 0.2 µm membrane.

### Library preparation and sequencing for ChIPseq

ChIPseq libraries were prepared from 5 ng of immunoprecipitated DNA using the NEBNext Ultra II DNA Library Prep Kit for Illumina (NEB, E7645), following the manufacturer’s instructions.

Briefly, 5 ng of ChIP DNA in 1× TE (50 µL) was subjected to end-repair and A-tailing (End Prep) and directly processed for adaptor ligation using a 10-fold–diluted NEBNext adaptor. Adaptor-ligated DNA was purified with a 0.9× SPRI bead cleanup (no size selection) and eluted in 15 µL 0.1× TE. Libraries were amplified with NEBNext Ultra II Q5 Master Mix and indexed primers using 7–8 PCR cycles, according to the recommended settings for 5 ng input. PCR products were purified with a 0.9× SPRI bead cleanup and eluted in 30 µL 0.1× TE. Final libraries were quantified by Qubit and their size distribution was assessed on an Agilent Bioanalyzer High Sensitivity DNA chip prior to pooling and Illumina sequencing.

### Bioinformatics analysis

#### RNAseq analysis

RNA-seq samples were sequenced using NovaSeq X plus sequencing device (paired-end, 150bp long reads, 86M to 284M de reads/sample) at Genewiz. The quality of each raw sequencing file (fastq) was verified with FastQC (https://www.bioinformatics.babraham.ac.uk/projects/fastqc/). Fastq were aligned the reference human genome (hg38) and counted on the Ensembl annotation from database (GRCh38.90) using STAR aligner (version 2.5.4a) ^34,35^.

Aligned BAM files were sorted and indexed using samtools (version 1.9)^35^ . Strand-specific, CPM-normalized coverage Bigwig files were generated using deepTools-bamCoverage (version 3.4.3) ^36^. For intron and exon-level quantification, reads were counted with Rsubread ::featureCounts() function (version 2.26.0) ^37^ against curated intron and exon annotations extracted with GTFtools ^38^ from a RefSeq-derived, Ensembl-formatted GTF.

Raw count matrices were filtered using HTSFilter ^39^ and filtered counts were then modeled using DESeq2 ^40^, with a design accounting for condition and replicate (∼ condition + replicate). Size factors, dispersion estimates, normalized counts, log2 Fold-Change, p-values and adjusted p-values were computed by DESeq2’s default pipeline.

Genes significantly differentially expressed in at least one contrast (adjusted p-value < 0.05, |log2 fold-change| > 0.7) were retained for clustering. Hierarchical clustering was performed on genes log2FC using Euclidean distance and complete linkageand visualized as heatmaps. Genes clusters were extracted by cutting the resulting dendrogram at a fixed height of 4 cutree() function

### HMM Readthrough analysis

Read-through transcripts detection was performed using the HMM pipeline as described (Jouffret et al., unpublished data). Read-through transcripts detected in at least one condition were retained for further analyses. Reads counting on RT and differential analysis was performed as described above using DESeq2’s default pipeline with the same normalization factors as used for introns.

### ChIPseq analysis

ChIP-seq samples were sequenced using NovaSeq X plus sequencing device (paired-end, 150bp long reads, 40M to 310M de reads/sample) at Genewiz. The quality of each raw sequencing fastq file was verified with FastQC (https://www.bioinformatics.babraham.ac.uk/projects/fastqc/). Adapter and low-quality bases were trimmed using Cutadapt ^41^. Trimmed reads were re-assessed with FastQC to confirm adapter/quality removal. Trimmed fastq were aligned custom combined human/Drosophila genome (hg38+dm6) using BWA-MEM^42^ .

Aligned BAM files were sorted, indexed and PCR duplicates were removed using samtools (version 1.9) ^35^[2]. Spike-in Drosophila Melanogaster reads were then filtered to generate a human-only BAM files. From these BAM, a coverage was computed with deeptools bamCoverage (version 3.4.3) ^36^ to produce strand specific bigwig output.

Matrices to draw metaprofiles were computed with deeptools computeMatrix (version 3.4.3) ^36^ giving one row per genes, bins of 50bp as columns (TSS-2kb;TSS ;TSS2kb) and median coverage as value.

For analysis of pol II occupancy at TSS , reads between -500 and +100 around TSS were counted for each genes using Rsubread ::featureCounts() function (version 2.26.0) ^37^. Background signal (corresponding to 40 reads/1kb, calculated as the median in the gene population of signal computed from -2000 to -1000 to each TSS) was then subtracted, and genes with a negative value in at least one sample were removed, resulting in values called “counts [TSS]”. Resulting count matrices were filtered using HTSFilter ^39^ and filtered counts were then modeled using DESeq2 ^40^, with a design accounting for condition and replicate (∼ condition + replicate). Size factors, dispersion estimates, normalized counts, log2 Fold-Change, p-values and adjusted p-values were computed by DESeq2’s default pipeline.

Genes with significantly different RNA pol II or S5-P RNA pol II occupancy at TSS in at least one contrast (adjusted p-value < 0.05, |log2 fold-change| > 1) were retained for clustering. Hierarchical clustering was performed on log2FC using Euclidean distance and complete linkage and visualized as heatmaps. Genes clusters were extracted by cutting the resulting dendrogram at a fixed height of 6 for RNA pol II and 8 for S5-P RNA pol II cutree() function.

For escape index calculation, reads between +500 and +2kb after each TSS were computed for each gene using Rsubread ::featureCounts() function (version 2.26.0)^37^ , background (corresponding to 40 reads/1kb, see above) was subtracted and genes with a negative value in at least one sample were removed, resulting in values called “counts [+500; +2000]. Escape index was calculating for each gene by The formula : (counts [+500;+2000]/1501/(counts [+500;+2000]/1501 + counts [TSS]/601).

## Reagents

### Antibodies

#### Primary antibodies

Mouse monoclonal Anti Flag-M2 antibody (F1824 Sigma), IF (1/1000)

Mouse Monoclonal Anti-HA.11 Epitope Tag Antibody anti-HA.11 - 16B12 (BioLegend), WB (1/5000)

Rabbit polyclonal Anti TRF1 antibody (homemade, generous gift from L. Crabbe -Vert), IF (1/200)

Rabbit monoclonal Anti-Rpb1 NTD D8L4Y (14958 Cell Signaling)

Rabbit monoclonal Anti-Phospho Rpb1 CTD (Ser5) (D9N51) (13523 Cell Signaling)

Rabbit monoclonal Anti Cas9 (8C1-F10) (61757 Active Motif)

### Secondary antibodies

Goat anti mouse Alexa Fluor red 594 (Thermo Fisher) coupled secondary antibody, IF (1/1000) Goat anti Rabbit Alexa Fluor 488 (Thermo Fisher) coupled secondary antibody, IF (1/1000) Goat anti mouse HRP coupled secondary antibody (Sigma), WB (1/10 000)

sgRNA sequences

OGG1: 5’-AAGGGTTCCAAAGGGCCGGA-3’

Telomere: 5’-TAGGGTTAGGGTTAGGGTTA-3’

Promoters :

CAV1

CAV1 1268 FW : TTTATGTTTTCCTAATGGAGAG

CAV1_1481 FW : GAGTCCTGGGGACAGTCCCC

CAV1 2015 FW : GATCTTAGATAAAGCTGGAA

CAV1 2322 FW : TTAAAAAGCTGGCTACGCGC

HAPLN1

HAPLN1 52 FW : CTTGAGAGCATCCGAACTCC HAPLN1

674 FW : AAACCCCAGAGCGGGATACT HAPLN1

794 FW : GCACATTAGGCTGTAATTAG

NCKAP5L

NCKAP5L -139 FW : GGCAGACTTAGCTCGCGTCC

NCKAP5L -12 FW : CCCTTTAAAGCCTTGACATC

NCKAP5L +136 FW : TTGAAAGGGCGGCGAGTGCG

NCKAP5L +163 FW : GATCTGTAGCTCCGGGCCGG

SLC41A1

SLC41A1 -287 FW : AGTCTGCTGGGGCTCCGACG

SLC41A1 -109 FW : GCCTCCTGGGCACGAAGCGA

SLC41A1 72 FW : TCCGCGGGCAAGAAGCGATT

Primers

RPLP0 int9

RPLP0 int9 FW : TGCTCTTCATGTTGCCCCAG

RPLP0 int9 REV : GTAGCCGGTGTGAGTAGACC

GAPDH int8

GAPDH int8 FW : TATGTGGCTGGGGCCAGAGA

GAPDH int8 REV : CAGGGCCCTTTTTCTGAGCC

CAV1 prom

CAV1 prom FW : TGGGGACTTTCGGGATTGTG

CAV1 prom REV: ACCCGTGGAATGTTCCCTT

CAV1 int2

CAV1 int2 FW : TCCCGGGATGAGTCAGAGAG

CAV1 int2 FW : TTGAGGATCCCCACACTTGC

HAPLN1 prom

HAPLN1 prom FW : CCTCCCTTCCTCCCTGAAGT

HAPLN1 prom REV : GCAAAGCGTCACCAGGAAAG

HAPLN1 int1

HAPLN1 int1 FW :GAGAGGGGTCATTAGCCTGC

HAPLN1 int1 REV : AACGGTTTGGGTTTGGAGGT

NCKAP5L prom

NCKAP5L Prom FW : GGAGGTCCTTCCCACCAAAG

NCKAP5L Prom REV : CTCCCTAGCAGCCAATCCAC

NCKAP5L int1

NCKAP5L int1 FW : TCTGCAGAAAAGGCCCTCTG

NCKAP5L int1 REV : GGCAGCCGCTTTCTTAGTTG

SLC41A1 prom

SLC41A1 P1 FW : GGTTCCCGTCACTCCTCCTA

SLC41A1 P1 REV : TCTCGGATGGACAACAGCAG

SLC41A1 int1

SLC41A1 int1 FW : CAGGAAGAGCGGGATGGTTT

SLC41A1 int1 REV : ATGGACTCCTGCTCAGTTGC

NF2 Read-Through

NF2 RT1 FW : ACCAGCAAAGGGGGAAACAA

NF2 RT1 REV: TGTGCGGCTCAGAATAGCAA

NF2 RT2 FW : AAAAGAACGCCAGCAGTGAC

NF2 RT2 REV : GGCCCTAGTGTGCCTAATGG

TPT1 Read-Through

TPT1 RT1 FW : GGCCATGAGAACCTTTGGGA

TPT1 RT1 REV : CCAGTTTGTGCTCCTTGCTC

TPT1 RT2 FW : CCTTCTCTACAGGCTTGGGC

TPT1 RT2 REV : GGGGAGGGAGACAGGATTCA

## Acknowledgments

We are grateful to Brigitte Schmidt and Bruce Armitage (Carnegie Mellon University) for providing MG2i dye. We would like to thank Xavier Renaudin for critical reading of the manuscript. We thank the LITC imaging facility at CBI.

## Author contributions

Luana Cintori (investigation, formal analysis, writing-original draft), Catherine Chailleux (investigation, formal analysis, data curation), Pauline Maller (investigation, formal analysis), Emma Lehir (investigation, formal analysis), Celine Reyes (investigation, formal analysis), Tomas Caetano (data curation, formal analysis), Virginie Jouffret (data curation, formal analysis), Marion Aguirrebengoa (data curation, formal analysis), Anne-Marie Di Guilmi (investigation, formal analysis), Capucine Siberchicot (investigation, formal analysis), Anna Campalans (investigation, supervision, funding acquisition, review and editing), Valérie Bergoglio (investigation, formal analysis, writing-original draft, writing-review and editing), Didier Trouche (supervision, conceptualization, funding acquisition, writing-original draft, writing-review&editing), Yvan Canitrot (investigation, conceptualization, supervision, funding acquisition, writing-original draft, writing-review&editing)

## Supplementary data

Supplementary data is available

## Data availability

## References

1. Kohen R, Nyska A. Oxidation of biological systems: oxidative stress phenomena, antioxidants, redox reactions, and methods for their quantification. Toxicol Pathol. 2002;30(6):620–50. doi:10.1080/01926230290166724

2. Tubbs A, Nussenzweig A. Endogenous DNA Damage as a Source of Genomic Instability in Cancer. Cell. Feb 09 2017;168(4):644–656. doi:10.1016/j.cell.2017.01.002

3. Caldecott KW. Mammalian DNA base excision repair: Dancing in the moonlight. DNA Repair (Amst). Sep 2020;93:102921. doi:10.1016/j.dnarep.2020.102921

4. David SS, O’Shea VL, Kundu S. Base-excision repair of oxidative DNA damage. Nature. Jun 21 2007;447(7147):941–50. doi:10.1038/nature05978

5. Heine GF, Horwitz AA, Parvin JD. Multiple mechanisms contribute to inhibit transcription in response to DNA damage. J Biol Chem. Apr 11 2008;283(15):9555–61. doi:10.1074/jbc.M707700200

6. Adamowicz M, Hailstone R, Demin AA, et al. XRCC1 protects transcription from toxic PARP1 activity during DNA base excision repair. Nat Cell Biol. Dec 2021;23(12):1287–1298. doi:10.1038/s41556-021-00792-w

7. Thomas QA, Wu L, Lesage E, et al. Oxidative stress triggers RNAPII arrest through PARylation and DNA damage. Nat Commun. Aug 10 2026;17(1)doi:10.1038/s41467-026-76443-1

8. Charlet-Berguerand N, Feuerhahn S, Kong SE, et al. RNA polymerase II bypass of oxidative DNA damage is regulated by transcription elongation factors. EMBO J. Nov 29 2006;25(23):5481–91. doi:10.1038/sj.emboj.7601403

9. Obermann T, Sakshaug T, Kanagaraj VV, et al. Genomic 8-oxoguanine modulates gene transcription independent of its repair by DNA glycosylases OGG1 and MUTYH. Redox Biol. Feb 2025;79:103461. doi:10.1016/j.redox.2024.103461

10. Giannakakis A, Zhang J, Jenjaroenpun P, et al. Contrasting expression patterns of coding and noncoding parts of the human genome upon oxidative stress. Sci Rep. May 29 2015;5:9737. doi:10.1038/srep09737

11. Vilborg A, Sabath N, Wiesel Y, et al. Comparative analysis reveals genomic features of stress-induced transcriptional readthrough. Proc Natl Acad Sci U S A. Oct 03 2017;114(40):E8362–E8371. doi:10.1073/pnas.1711120114

12. Endutkin AV, Dvornikova AP, Zharkov DO. 8-Oxoguanine: A Lesion, an Epigenetic Mark, or a Molecular Signal? Int J Mol Sci. Dec 06 2025;26(24)doi:10.3390/ijms262411799

13. Fleming AM, Ding Y, Burrows CJ. Oxidative DNA damage is epigenetic by regulating gene transcription via base excision repair. Proc Natl Acad Sci U S A. Mar 07 2017;114(10):2604–2609. doi:10.1073/pnas.1619809114

14. Cogoi S, Ferino A, Miglietta G, Pedersen EB, Xodo LE. The regulatory G4 motif of the Kirsten ras (KRAS) gene is sensitive to guanine oxidation: implications on transcription. Nucleic Acids Res. Jan 25 2018;46(2):661–676. doi:10.1093/nar/gkx1142

15. Pan L, Zhu B, Hao W, et al. Oxidized Guanine Base Lesions Function in 8-Oxoguanine DNA Glycosylase-1-mediated Epigenetic Regulation of Nuclear Factor κB-driven Gene Expression. J Biol Chem. Dec 02 2016;291(49):25553–25566. doi:10.1074/jbc.M116.751453

16. Tanushi X, Pinna G, Vandamme M, et al. OGG1 competitive inhibitors show important off-target effects by directly inhibiting efflux pumps and disturbing mitotic progression. Front Cell Dev Biol. 2023;11:1124960. doi:10.3389/fcell.2023.1124960

17. Lebraud E, Pinna G, Siberchicot C, et al. Chromatin recruitment of OGG1 requires cohesin and mediator and is essential for efficient 8-oxoG removal. Nucleic Acids Res. Sep 18 2020;48(16):9082–9097. doi:10.1093/nar/gkaa611

18. Amouroux R, Campalans A, Epe B, Radicella JP. Oxidative stress triggers the preferential assembly of base excision repair complexes on open chromatin regions. Nucleic Acids Res. May 2010;38(9):2878–90. doi:10.1093/nar/gkp1247

19. Ballmaier D, Epe B. DNA damage by bromate: mechanism and consequences. Toxicology. Apr 17 2006;221(2-3):166–71. doi:10.1016/j.tox.2006.01.009

20. Kawanishi S, Murata M. Mechanism of DNA damage induced by bromate differs from general types of oxidative stress. Toxicology. Apr 17 2006;221(2-3):172–8. doi:10.1016/j.tox.2006.01.002

21. Roth SJ, Heinz S, Benner C. ARTDeco: automatic readthrough transcription detection. BMC Bioinformatics. May 26 2020;21(1):214. doi:10.1186/s12859-020-03551-0

22. Rutkowski AJ, Erhard F, L’Hernault A, et al. Widespread disruption of host transcription termination in HSV-1 infection. Nat Commun. May 20 2015;6:7126. doi:10.1038/ncomms8126

23. Sandoval J, Rodríguez JL, Tur G, et al. RNAPol-ChIP: a novel application of chromatin immunoprecipitation to the analysis of real-time gene transcription. Nucleic Acids Res. 2004;32(11):e88. doi:10.1093/nar/gnh091

24. Pan L, Hao W, Zheng X, et al. OGG1-DNA interactions facilitate NF-κB binding to DNA targets. Sci Rep. Mar 07 2017;7:43297. doi:10.1038/srep43297

25. Barnes RP, de Rosa M, Thosar SA, et al. Telomeric 8-oxo-guanine drives rapid premature senescence in the absence of telomere shortening. Nat Struct Mol Biol. Jul 2022;29(7):639–652. doi:10.1038/s41594-022-00790-y

26. Fouquerel E, Barnes RP, Uttam S, Watkins SC, Bruchez MP, Opresko PL. Targeted and Persistent 8-Oxoguanine Base Damage at Telomeres Promotes Telomere Loss and Crisis. Mol Cell. Jul 11 2019;75(1):117–130.e6. doi:10.1016/j.molcel.2019.04.024

27. Telmer CA, Verma R, Teng H, Andreko S, Law L, Bruchez MP. Rapid, specific, no-wash, far-red fluorogen activation in subcellular compartments by targeted fluorogen activating proteins. ACS Chem Biol. May 15 2015;10(5):1239–46. doi:10.1021/cb500957k

28. He J, Wang Y, Missinato MA, et al. A genetically targetable near-infrared photosensitizer. Nat Methods. Mar 2016;13(3):263–8. doi:10.1038/nmeth.3735

29. Vilborg A, Passarelli MC, Yario TA, Tycowski KT, Steitz JA. Widespread Inducible Transcription Downstream of Human Genes. Mol Cell. Aug 06 2015;59(3):449–61. doi:10.1016/j.molcel.2015.06.016

30. Hanna BMF, Michel M, Helleday T, Mortusewicz O. NEIL1 and NEIL2 Are Recruited as Potential Backup for OGG1 upon OGG1 Depletion or Inhibition by TH5487. Int J Mol Sci. Apr 27 2021;22(9)doi:10.3390/ijms22094542

31. Morland I, Rolseth V, Luna L, Rognes T, Bjørås M, Seeberg E. Human DNA glycosylases of the bacterial Fpg/MutM superfamily: an alternative pathway for the repair of 8-oxoguanine and other oxidation products in DNA. Nucleic Acids Res. Nov 15 2002;30(22):4926–36. doi:10.1093/nar/gkf618

32. Visnes T, Cázares-Körner A, Hao W, et al. Small-molecule inhibitor of OGG1 suppresses proinflammatory gene expression and inflammation. Science. Nov 16 2018;362(6416):834–839. doi:10.1126/science.aar8048

33. Li J, Li L, Tan Y, et al. Dynamic genome-wide mapping reveals how chromatin context shapes OGG1-mediated repair and related mutagenesis in human cells. Nucleic Acids Res. Nov 26 2025;53(22)doi:10.1093/nar/gkaf1330

34. Dobin A, Davis CA, Schlesinger F, et al. STAR: ultrafast universal RNA-seq aligner. Bioinformatics. Jan 01 2013;29(1):15–21. doi:10.1093/bioinformatics/bts635

35. Li H, Handsaker B, Wysoker A, et al. The Sequence Alignment/Map format and SAMtools. Bioinformatics. Aug 15 2009;25(16):2078–9. doi:10.1093/bioinformatics/btp352

36. Ramírez F, Ryan DP, Grüning B, et al. deepTools2: a next generation web server for deep-sequencing data analysis. Nucleic Acids Res. Jul 08 2016;44(W1):W160–5. doi:10.1093/nar/gkw257

37. Liao Y, Smyth GK, Shi W. The R package Rsubread is easier, faster, cheaper and better for alignment and quantification of RNA sequencing reads. Nucleic Acids Res. May 07 2019;47(8):e47. doi:10.1093/nar/gkz114

38. Li HD, Lin CX, Zheng J. GTFtools: a software package for analyzing various features of gene models. Bioinformatics. Oct 14 2022;38(20):4806–4808. doi:10.1093/bioinformatics/btac561

39. Rau A, Gallopin M, Celeux G, Jaffrézic F. Data-based filtering for replicated high-throughput transcriptome sequencing experiments. Bioinformatics. Sep 01 2013;29(17):2146–52. doi:10.1093/bioinformatics/btt350

40. Love MI, Huber W, Anders S. Moderated estimation of fold change and dispersion for RNA-seq data with DESeq2. Genome Biol. 2014;15(12):550. doi:10.1186/s13059-014-0550-8

41. Martin M. cutadapt removes adaptersequences from high-throughput sequencing reads. EMBnet; 2011.

42. Li H. Aligning sequence reads, clone sequences and assembly contigs with BWA-MEM. arXiv; 2013.

