## supplemental figures 1-6 for "Direct effects on gene transcription revealed by global and CRISPR-targeted oxidative DNA damage"

A

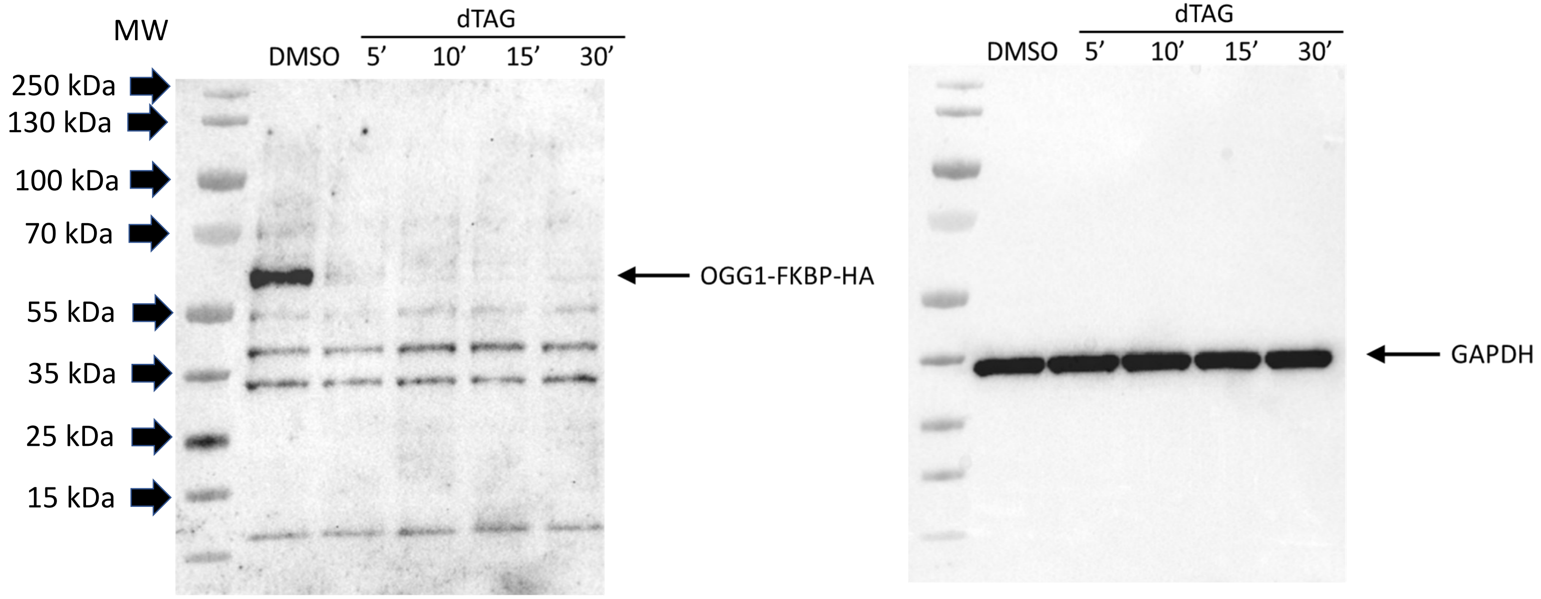


B

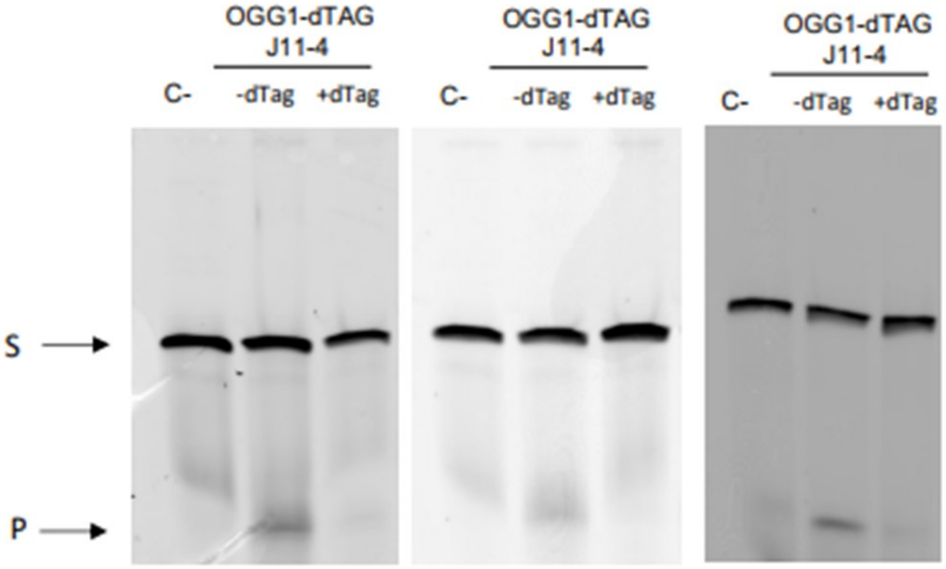


Figure Supp 1 Cintori et al.

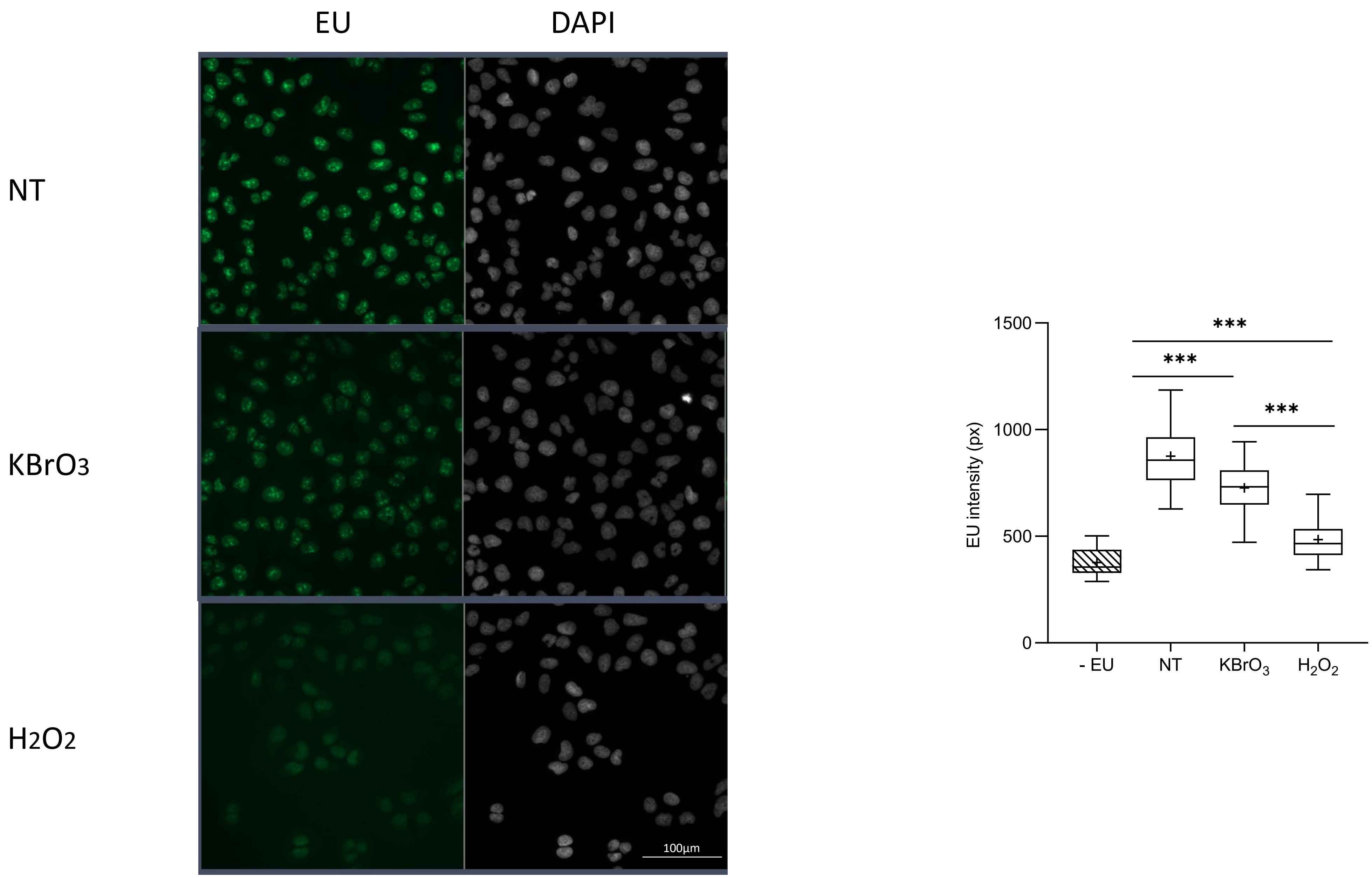


Supp Figure 2 Cintori et al.

### Gene Ontology

A

#### EnrichR_Molecular Function_ Cluster 2

| Index | Name | P-value | Adjusted p-value | Odds Ratio | Combined score |
|---|---|---|---|---|---|
| 1 | Gamma-aminobutyric Acid Transmembrane Transporter Activity (GO:0015185) | 0.001943 | 0.1587 | 38.31 | 239.17 |
| 2 | RNA Polymerase II-specific DNA-binding Transcription Factor Binding (GO:0061629) | 0.002058 | 0.1587 | 4.19 | 25.95 |
| 3 | Tat Protein Binding (GO:0030957) | 0.002417 | 0.1587 | 33.52 | 201.95 |
| 4 | Chloride Channel Regulator Activity (GO:0017081) | 0.004128 | 0.2033 | 24.37 | 133.80 |
| 5 | Protein Phosphatase 1 Binding (GO:0008157) | 0.01069 | 0.2806 | 14.10 | 64.02 |
| 6 | Organic Acid:Sodium Symporter Activity (GO:0005343) | 0.01497 | 0.2806 | 11.65 | 48.95 |
| 7 | Metal Ion Transmembrane Transporter Activity (GO:0046873) | 0.01987 | 0.2806 | 9.92 | 38.88 |
| 8 | SnoRNA Binding (GO:0030515) | 0.02119 | 0.2806 | 9.57 | 36.87 |
| 9 | DNA-binding Transcription Factor Binding (GO:0140297) | 0.02508 | 0.2806 | 2.79 | 10.29 |
| 10 | 3'-5'-RNA Exonuclease Activity (GO:0000175) | 0.02828 | 0.2806 | 8.12 | 28.93 |

#### EnrichR_Biological Process_ Cluster 2

| Index | Name | P-value | Adjusted p-value | Odds Ratio | Combined score |
|---|---|---|---|---|---|
| 1 | Cholesterol Biosynthetic Process (GO:0006695) | 8.238e-8 | 0.00003513 | 34.42 | 561.46 |
| 2 | Secondary Alcohol Biosynthetic Process (GO:1902653) | 8.238e-8 | 0.00003513 | 34.42 | 561.46 |
| 3 | Sterol Biosynthetic Process (GO:0016126) | 1.242e-7 | 0.00003531 | 31.77 | 505.18 |
| 4 | Cholesterol Metabolic Process (GO:0008203) | 0.00002284 | 0.004871 | 11.77 | 125.83 |
| 5 | Negative Regulation of Endoplasmic Reticulum Unfolded Protein Response (GO:1900102) | 0.0003669 | 0.06259 | 25.30 | 200.12 |
| 6 | Glutathione Transport (GO:0034635) | 0.0005505 | 0.07788 | 89.40 | 670.92 |
| 7 | Steroid Biosynthetic Process (GO:0006694) | 0.0007455 | 0.07788 | 19.27 | 138.77 |
| 8 | Positive Regulation of Aggrephagy (GO:1905337) | 0.0008217 | 0.07788 | 67.05 | 476.31 |
| 9 | Gamma-aminobutyric Acid Import (GO:0051939) | 0.0008217 | 0.07788 | 67.05 | 476.31 |
| 10 | Response to UV-C (GO:0010225) | 0.001145 | 0.09765 | 53.64 | 363.25 |

B

#### EnrichR_Molecular Function_ Cluster3

| Index | Name | P-value | Adjusted p-value | Odds Ratio | Combined score |
|---|---|---|---|---|---|
| 1 | RNA Polymerase II Transcription Regulatory Region Sequence-Specific DNA Binding (GO:0000977) | 0.0005337 | 0.09172 | 2.15 | 16.18 |
| 2 | Cis-regulatory Region Sequence-Specific DNA Binding (GO:0000987) | 0.001193 | 0.09172 | 2.15 | 14.48 |
| 3 | RNA Polymerase II Cis-Regulatory Region Sequence-Specific DNA Binding (GO:0000978) | 0.001555 | 0.09172 | 2.10 | 13.61 |
| 4 | UDP-glucosyltransferase Activity (GO:0035251) | 0.007008 | 0.3101 | 18.66 | 92.55 |
| 5 | Aspartic-type Peptidase Activity (GO:0070001) | 0.01558 | 0.3515 | 11.66 | 48.52 |
| 6 | S-adenosylmethionine-dependent Methyltransferase Activity (GO:0008757) | 0.02423 | 0.3515 | 5.10 | 18.98 |
| 7 | Transcription Cis-Regulatory Region Binding (GO:0000976) | 0.02907 | 0.3515 | 1.93 | 6.84 |
| 8 | DNA-binding Transcription Repressor Activity, RNA Polymerase II-specific (GO:0001227) | 0.03902 | 0.3515 | 2.50 | 8.10 |
| 9 | Mitochondrial Large Ribosomal Subunit Binding (GO:0140978) | 0.05237 | 0.3515 | 23.22 | 68.48 |
| 10 | N-acetyl-beta-D-glucosaminide Beta-(1,3)-Galactosyltransferase Activity (GO:0008499) | 0.05237 | 0.3515 | 23.22 | 68.48 |

#### EnrichR_Biological Process_ Cluster3

| Index | Name | P-value | Adjusted p-value | Odds Ratio | Combined score |
|---|---|---|---|---|---|
| 1 | Regulation of Apoptotic Process (GO:0042981) | 0.9967 | 0.9967 | 0.25 | 0.00 |
| 2 | Protein Metabolic Process (GO:0019538) | 0.9944 | 0.9955 | 0.27 | 0.00 |
| 3 | DNA Damage Response (GO:0006974) | 0.9936 | 0.9955 | 0.20 | 0.00 |
| 4 | Protein Modification Process (GO:0036211) | 0.9916 | 0.9949 | 0.21 | 0.00 |
| 5 | Proteasome-mediated Ubiquitin-Dependent Protein Catabolic Process (GO:0043161) | 0.9896 | 0.9940 | 0.22 | 0.00 |
| 6 | Negative Regulation of Programmed Cell Death (GO:0043069) | 0.9884 | 0.9939 | 0.22 | 0.00 |
| 7 | Positive Regulation of RNA Biosynthetic Process (GO:1902680) | 0.9842 | 0.9908 | 0.32 | 0.01 |
| 8 | Chromatin Organization (GO:0006325) | 0.9804 | 0.9880 | 0.25 | 0.01 |
| 9 | Positive Regulation of Cytokine Production (GO:0001819) | 0.9795 | 0.9880 | 0.26 | 0.01 |
| 10 | Nervous System Development (GO:0007399) | 0.9789 | 0.9880 | 0.34 | 0.01 |

Supp Figure 3 Cintori et al.

C

### Gene Ontology

#### EnrichR_Molecular Function_ Cluster 4

| Index | Name | P-value | Adjusted p-value | Odds Ratio | Combined score |
|---|---|---|---|---|---|
| 1 | Outward Rectifier Potassium Channel Activity (GO:0015271) | 0.0005515 | 0.1257 | 21.60 | 162.04 |
| 2 | Potassium Channel Activity (GO:0005267) | 0.002946 | 0.2342 | 7.23 | 42.12 |
| 3 | Voltage-gated Potassium Channel Activity (GO:0005249) | 0.003081 | 0.2342 | 7.13 | 41.25 |
| 4 | Potassium Ion Leak Channel Activity (GO:0022841) | 0.006098 | 0.2458 | 19.41 | 98.99 |
| 5 | Transmembrane Receptor Protein Tyrosine Kinase Activity (GO:0004714) | 0.006733 | 0.2458 | 8.36 | 41.81 |
| 6 | Leak Channel Activity (GO:0022840) | 0.009470 | 0.2458 | 15.09 | 70.33 |
| 7 | Glutamate Receptor Binding (GO:0035254) | 0.01460 | 0.2458 | 11.81 | 49.92 |
| 8 | Notch Binding (GO:0005112) | 0.01460 | 0.2458 | 11.81 | 49.92 |
| 9 | Protein Heterodimerization Activity (GO:0046982) | 0.02049 | 0.2458 | 3.33 | 12.96 |
| 10 | G Protein-Coupled Receptor Binding (GO:0001664) | 0.02280 | 0.2458 | 3.88 | 14.68 |

#### EnrichR_Biological Process_ Cluster4

| Index | Name | P-value | Adjusted p-value | Odds Ratio | Combined score |
|---|---|---|---|---|---|
| 1 | Signal Transduction by P53 Class Mediator (GO:0072331) | 0.00007526 | 0.04913 | 12.82 | 121.71 |
| 2 | Cellular Response to Acid Chemical (GO:0071229) | 0.0001573 | 0.04913 | 16.68 | 146.10 |
| 3 | Response to Amino Acid (GO:0043200) | 0.0001936 | 0.04913 | 15.73 | 134.47 |
| 4 | Intrinsic Apoptotic Signaling Pathway by P53 Class Mediator (GO:0072332) | 0.0001936 | 0.04913 | 15.73 | 134.47 |
| 5 | Cellular Response to Amino Acid Stimulus (GO:0071230) | 0.0004010 | 0.06982 | 12.80 | 100.09 |
| 6 | Amino Sugar Metabolic Process (GO:0006040) | 0.0004127 | 0.06982 | 24.14 | 188.12 |
| 7 | Cellular Response to Starvation (GO:0009267) | 0.0009482 | 0.1037 | 5.70 | 39.70 |
| 8 | Regulation of Aldosterone Biosynthetic Process (GO:0032347) | 0.001115 | 0.1037 | 54.38 | 369.71 |
| 9 | Cardiac Cell Fate Commitment (GO:0060911) | 0.001115 | 0.1037 | 54.38 | 369.71 |
| 10 | Protein Repair (GO:0030091) | 0.001115 | 0.1037 | 54.38 | 369.71 |

Supp Figure 3 Cintori et al.

A

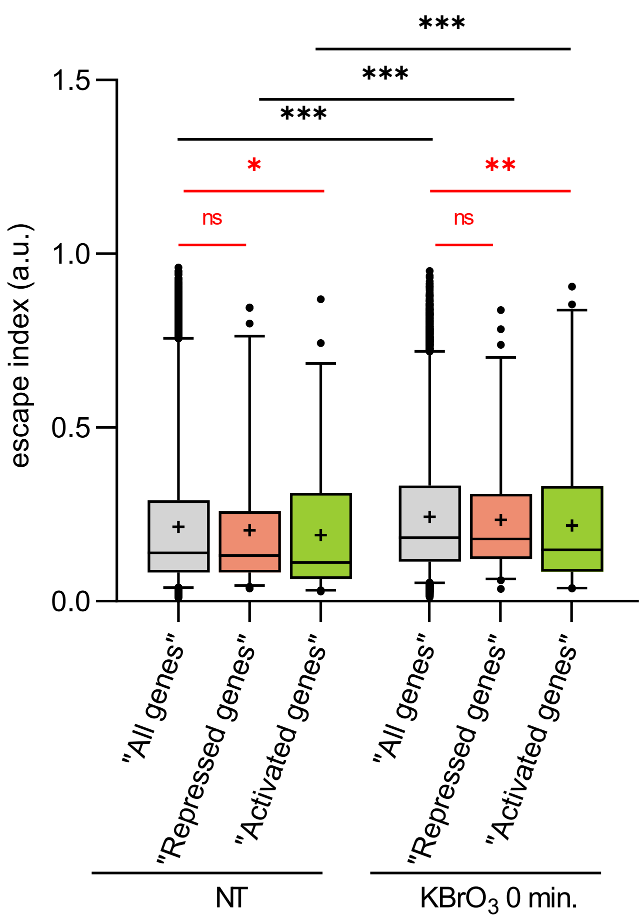


B

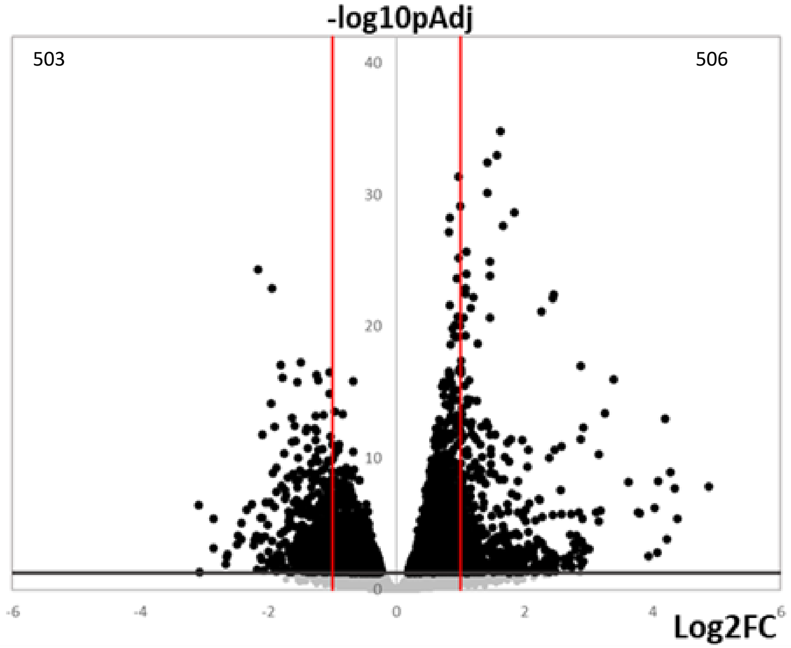


Supp Figure 4 Cintori et al.

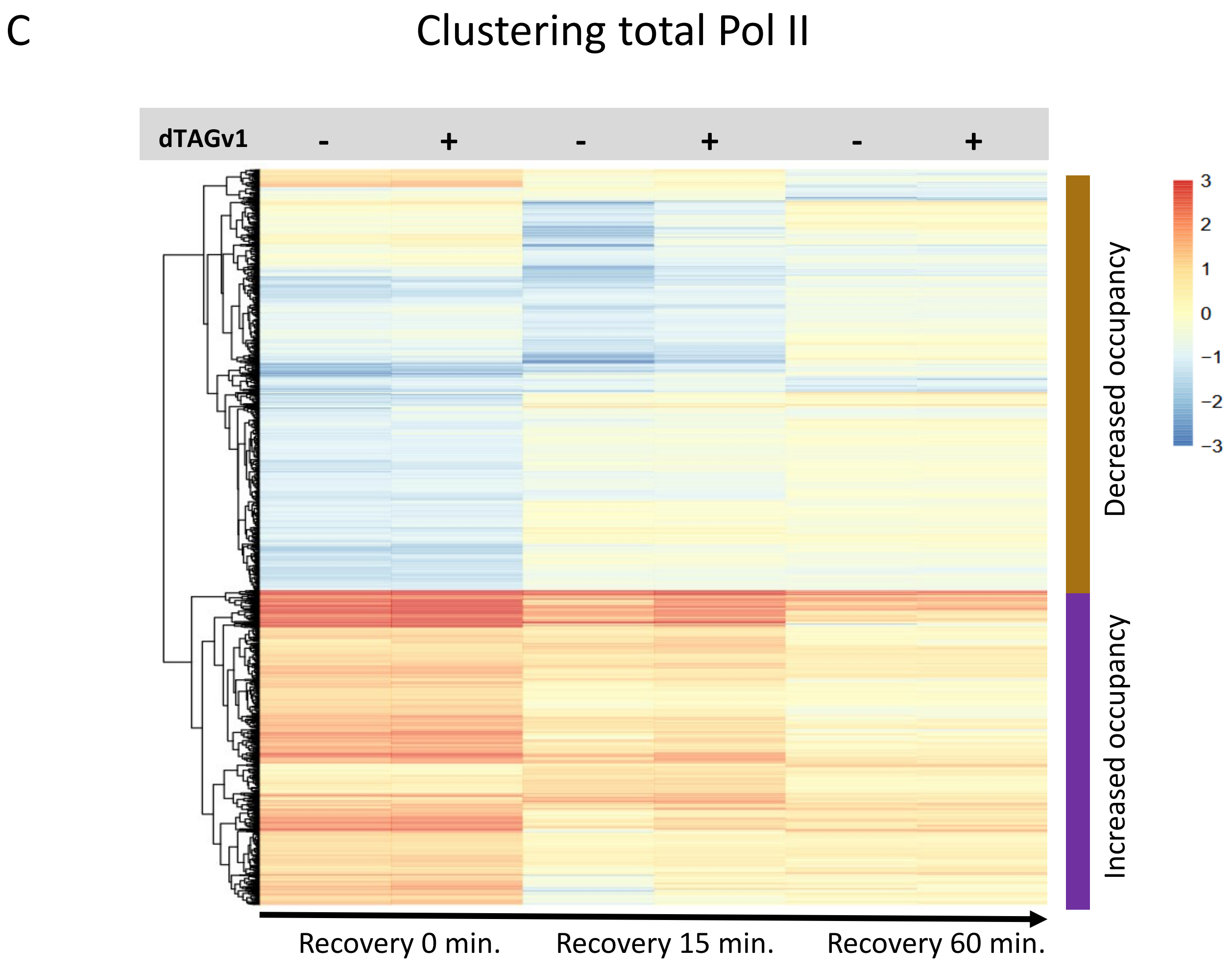


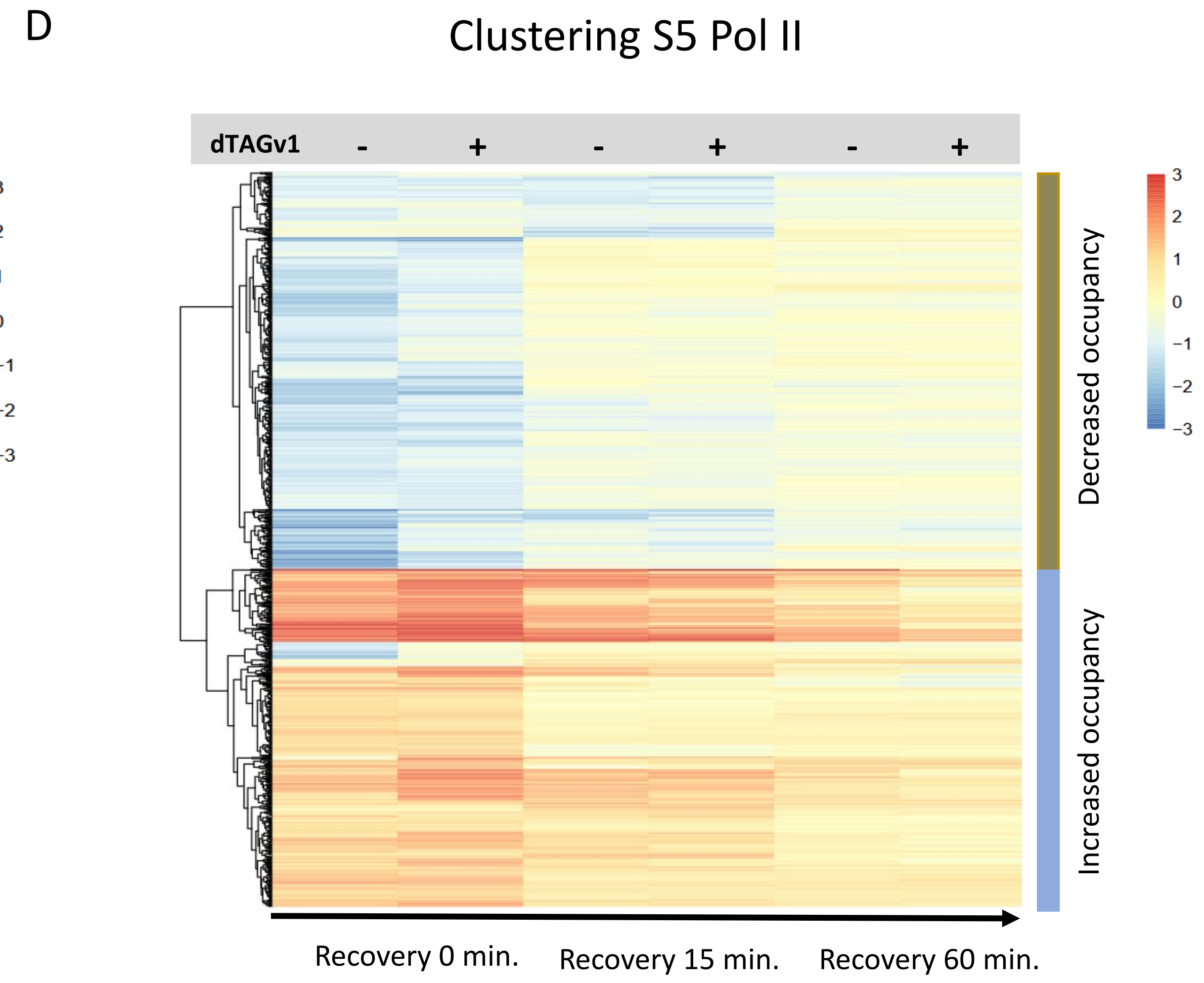


Supp Figure 5 Cintori et al.

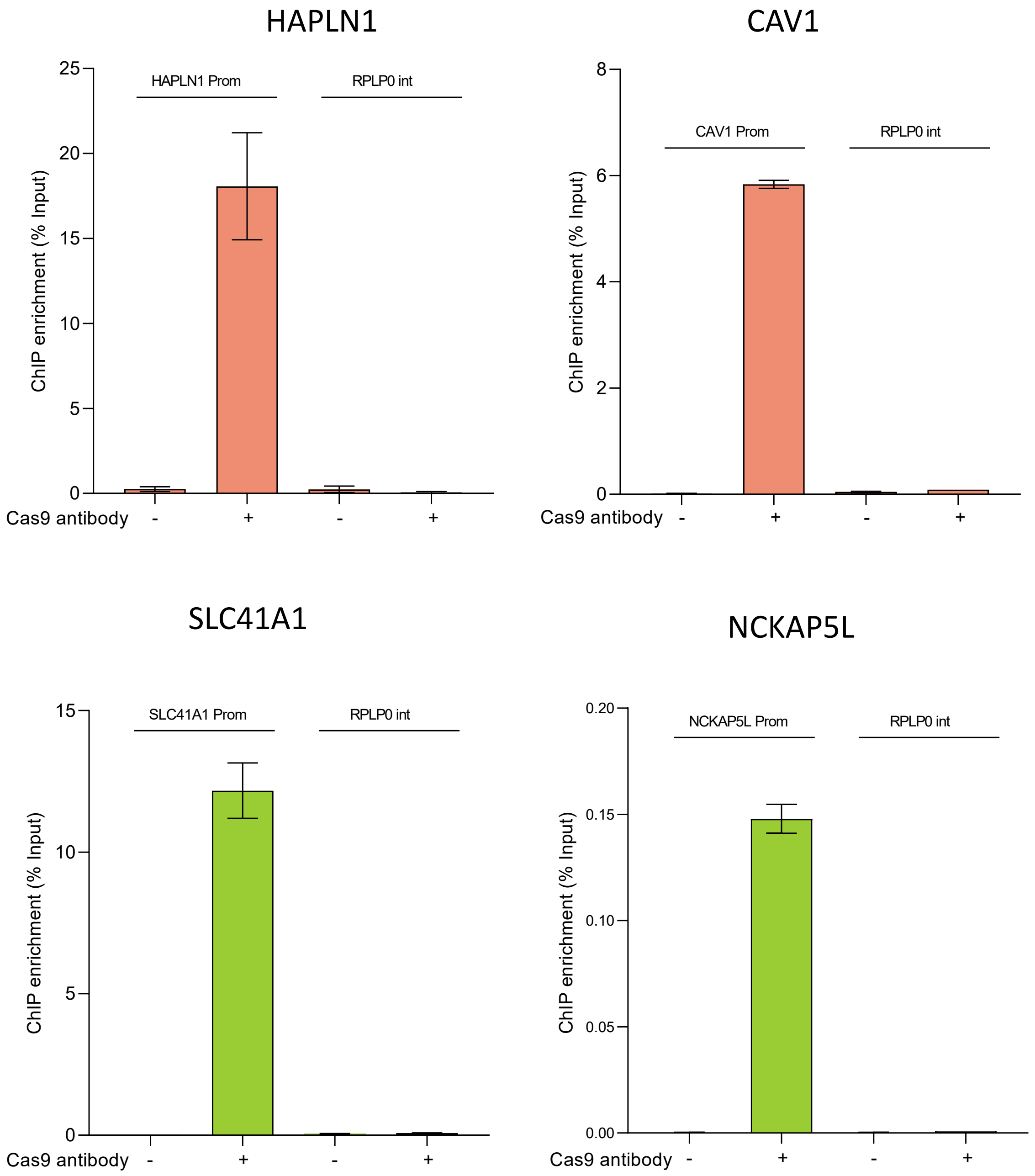


Supp Figure 6 Cintori et al.
