## Supplementary material for "Direct effects on gene transcription revealed by global and CRISPR-targeted oxidative DNA damage": legends of supplemental figures

**Legend to supplementary figures**

**Figure S1 :**

A. Full western blot image used in Figure 1. **B.** Replicate experiments from Figure 1C.

**Figure S2:**

Representative images of EU incorporation during 30 min of KBr0_3_ 40mM or H_2_O_2_ 1mM treatment compared to untreated condition. Nuclei are counterstained with DAPI. Images were acquired with a 20X objective with the same exposure time. Quantification of EU incorporation in the different conditions of treatment.

**Figure S3 :**

Gene Ontology analysis (<https://maayanlab.cloud/Enrichr>. ) of cluster 2 (**A**), cluster 3 (**B**) and cluster 4 (**C**). The first 10 GO terms from “Molecular Function” and “Biological processes” are shown.

**Figure S4 :**

**A.** Replicate experiment for Figure 6B. **B.** Volcano plot showing RNA PolII recruitment at promoter regions after KBrO3 treatment at time 0 compared with untreated control. On the upper corner left and right are the number of genes with -log10pAdj>1,3 and Log2FC<-1 or >1. **C.** Genes with a padj<0.05 and a Log2FC<-1 or >1 in at least one condition from the differential analysis of total pol II occupancy at TSS described in Figure 6A were subjected to unsupervised clustering based on their Log2FC. This led to the identification of two groups of genes, one harbouring a decreased RNA pol II occupancy, and one experiencing an increase of RNA pol II occupancy. These two clusters were used for Figure 6C. **D.** Same as in C, except that S5-P pol II occupancy was analysed. Two groups of genes were identified, one harbouring a decreased S5-P RNA pol II occupancy, and one experiencing an increase of S5-P RNA pol II occupancy. These two clusters were used for Figure 6D.

**Figure S5:**

FAP-dCas9 cells were transfected with the indicated sgRNA. 24 hours later, they were harvested and subjected to a ChIP experiment using anti-Cas9 antibody. The presence of the targeted promoter as well as a control DNA sequence (RPLP0) were analysed for each sgRNAs by qPCR. A representative experiment out of two is shown.

Figure S6.
